# Spontaneous Transmembrane Pore Formation by Short-chain Synthetic Peptide

**DOI:** 10.1101/2021.02.12.430925

**Authors:** Jaya Krishna Koneru, Dube Dheeraj Prakashchand, Namita Dube, Pushpita Ghosh, Jagannath Mondal

**Affiliations:** Tata Institute of Fundamental Research, Center for Interdisciplinary sciences, Hyderabad, Telangana 500046, India

## Abstract

Amphiphilic *β*-peptides, which are synthetically designed short-chain helical foldamer of *β*-amino acids, are established potent biomimetic alternatives of natural antimicrobial peptides. An intriguing question is: how does the distinct molecular architecture of these short-chain and rigid synthetic peptides translates to its potent membrane disruption ability? Here, we address this question via a combination of all atom and coarse-grained molecular dynamics simulations of the interaction of mixed phospholipid bilayer with an antimicrobial 10-residue globally amphiphilic helical *β*-peptide at wide range of concentrations. The simulation demonstrates that multiple copies of this synthetic peptide, initially placed in aqueous solution, readily self-assemble and adsorb at membrane interface. Subsequently, beyond a threshold peptide-to-lipid ratio, the surface-adsorbed oligomeric aggregate moves inside the membrane and spontaneously forms stable water-filled transmembrane pore via a cooperative mechanism. The defects induced by these pores lead to the dislocation of interfacial lipid head groups, membrane thinning and substantial water leakage inside the hydrophobic core of the membrane. A molecular analysis reveals that, despite having a short architecture, these synthetic peptides, once inside the membrane, would stretch themselves towards the distal leaflet in favour of potential contact with polar head groups and interfacial water layer. The pore formed in coarse-grained simulation was found to be resilient upon structural refinement. Interestingly, the pore-inducing ability was found to be elusive in a non-globally amphiphilic sequence isomer of the same *β*-peptide, indicating strong sequence dependence. Taken together, this work put forward key perspectives of membrane-activity of minimally designed synthetic biomimetic oligomers relative to the natural antimicrobial peptides.

**STATEMENT OF SIGNIFICANCE:** The development of bacterial resistance to conventional antibiotics is a major concern towards public health. Antimicrobial peptides, which provide a natural defence against a large range of pathogens, including bacteria and fungi, are emerging as a sustainable substitute of antibiotics. However, serious issues with the naturally occurring antimicrobial peptides which have prevented their wide-spread appreciations are their susceptibility to degradation and lack of specificity for microbial targets. In this regard, synthetic biomimetic peptides are coming up as a viable alternative. In this work we provide clarity on how these synthetic antimicrobial peptides, which often involves distinctly short architecture, acts on the membrane. We show that despite its short architecture, a 10-residue biomimetic peptide, *β*-peptide, can spontaneously form stable membrane-spanning pore and induce water-leakage inside the membrane.

## 1 INTRODUCTION

The development of bacterial resistance to conventional antibiotics is a major concern towards public health. Antimicrobial peptides, which provide a natural defence against a large range of pathogens, including bacteria and fungi, are emerging as a sustainable substitute of antibiotics. The peptide-based antimicrobial agents are less prone to bacterial resistance than the natural antimicrobial agents and have got well-deserved prominence(1–4). However, serious issues with the naturally occurring antimicrobial peptides which have prevented their wide-spread appreciations are their susceptibility to degradation by cellular proteases and lack of specificity for microbial targets over host cells. In this regard, synthetic oligomers and synthetic peptides which adopt helical conformations, are coming up as a viable alternative to overcome these limitations of natural antimicrobial peptides(5–8). The present work focuses on a promising class of potentially novel biomimetic anti-microbial synthetic peptides, referred as *β*-peptides and provides key molecular insights into the underlying mechanisms of their membrane-disruption capability.

*β*-peptides are synthetic oligomers of *β*-amino acids and are rationally designed to mimic and improve the biological activities of their natural counter parts, namely *α*-peptides. The backbone of a *β*-amino acid contains an additional carbon atom compared to that of *α*-amino acid, which provides an additional site for introducing side chain and makes the *β*-peptides resistant to degradation by natural proteases(9). Synthetic oligomers and random copolymers of *β* amino acids(10, 11) have shown potent antibacterial(12–14) and antifungal (15, 16) activity. The synthetic peptide of our current interest is an amphiphilic 10-residue-length *β*-peptide sequence *β*Y-(ACHC-ACHC-*β*K)_3_ (hereby called ‘AAK’) where *β*Y, *β*K and ACHC refer to *β*-tyrosine, *β*-lysine and trans-2-amino cyclohexyl carboxylic acid (see Figure 1). The cyclic constraint of ACHC residues confers a stable 14-helical conformation to this significantly short *β*-peptide sequence.

**Figure 1:**
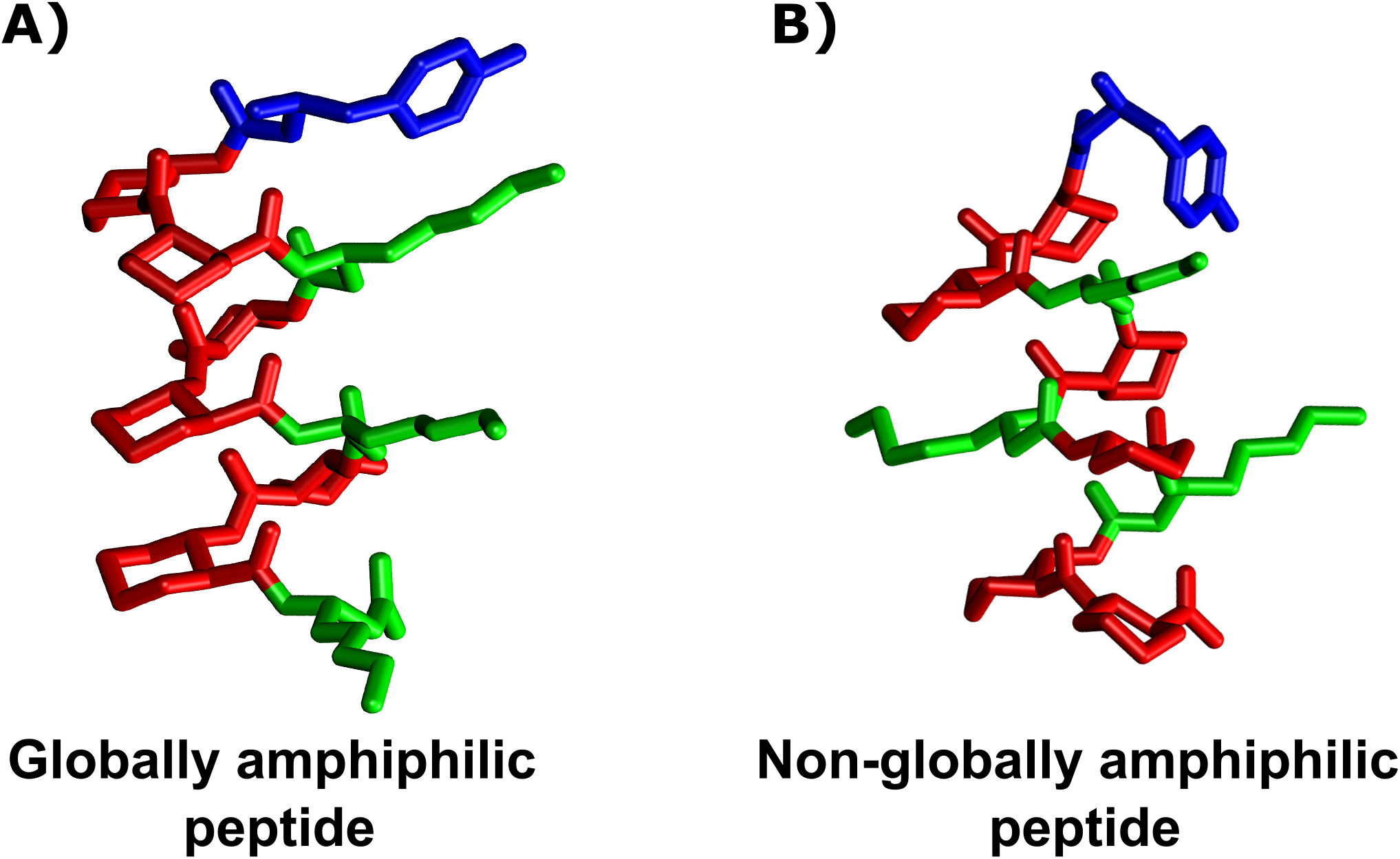
Three-dimensional representation of the *β*-peptides of our interest (‘AAK’). A: Globally amphiphilic (GA) and B) non-globally amphiphilic (non-GA) isomers of AAK. The *β*-peptide is shown in licorice representation with hydrophobic ACHC residues in red, *β*-lysine groups in green and *β*-tyrosines are in blue.

The ability to form stable 14-helical amphiphilic conformation by only ten *β*-amino acid residues provides the peptide a greater synthetic advantage over relatively long-chain naturally occurring antimicrobial peptides such as magainin, melittin or LL37. More importantly, Gellman and coworkers(12) have explored the interplay of sequence-specificity and the antimicrobial activity in AAK. Specifically, the design of a sequence-arrangement of AAK in which the hydrophobic ACHC and hydrophilic *β*-homolysine residues are globally segregated, gave rise to a globally amphiphilic (hereby called ‘GA’ isomer) sequence isomer (see Figure 1A)). Experimental investigations led by Gellman and coworkers(12) have previously shown that the GA isomer of AAK can impart strong anti-bacterial activity on both gram-positive and gram-negative bacteria at a very low minimum inhibitory concentration (MIC). The GA isomer of AAK were found to retain anti-fungal activity against *Candida albicans*(15) with MIC value ranging between 16-32 *µ*g/ml. AAK was also found to induce leakage of enzyme *β*-galactosidase from *Bacillus subtilis*(13), thereby suggesting membrane permeabilization by *β*-peptides. On a related study, Gellman group also had investigated the role of amphiphilicity in the antimicrobial activity of AAK by synthetically designing a non-globally amphiphilic sequence-isomer of AAK (hereby called ‘non GA’ isomer of AAK) via evenly distributing single hydrophilic *β*-homolysine residue on all faces of 14-helical AAK. It was found that the antibacterial potency of non GA isomer is significantly low, in contrast to its potent counterpart.(12). We also note that these two same sequences had later been investigated by Karlsson and coworkers group for their anti fungal activity(16).The lack of anti fungal activity in non-GA isomer was evident in that investigation. These observations suggested that the antimicrobial activity of AAK is strongly sequence-dependent, with GA isomer of AAK showing strong antibacterial and anti fungal propensity than its non GA counterpart. However, in all these experimental investigations, the relative comparison of minimum Inhibitory concentration (MIC) has remained the key metric of these two peptides’ antimicrobial activities. There is no report of more quantitative investigation of membrane affinity/cell leakage for these two *β*-peptide sequences. Together, while the precedent experimental investigations provide convincing macroscopic evidence of antimicrobial activity of AAK and the associated sequence-dependence, to the best of our knowledge, a molecule-level insight towards plausible mechanism of the antimicrobial activity of this ‘short-chain’ *β*-peptide, in comparison to its natural counterpart, has largely remained unexplored.

Since the origin of antimicrobial peptides’ activity is believed to be rooted in their interaction with the membrane of the bacteria(17), we wanted to explore how a membrane environment would respond to exposure to a growingly increasing concentration of *β*-peptides. In this current work, the pertinent question we ask: What is the plausible mechanism of membrane-disruption ability by these AAK ? Multiple hypotheses exist(1–3) in general to describe the principle steps leading to the membrane-disruption by antimicrobial peptides. Central to all prevailing hypotheses is the possibility of trans-membrane pore formation by antimicrobial peptides (18, 19). Does it form any transmembrane water channel? Or does it disrupt the membrane integrity in so-called “carpet like” (3, 4) mechanism? In fact, natural antimicrobial peptides namely magainin-H2, melittin are believed to form trans-membrane pore across the membrane bilayer(20, 21). However, it is also believed that in general, there is no unique mechanism and it strongly depends on the specific antimicrobial peptides of interest (22, 23). The question of membrane-disruption mechanism is even more relevant for AAK mainly due to the significantly distinct molecular architecture and *short* sequence lengths of these rigid *β*-peptides compared to its flexible and sequentially long large molecular weights natural counterparts like magainin-H2 and melittin.

In the current work, we address these questions via coarse-grained molecular dynamics (MD) simulations of interactions of GA and non GA isomers AAK with a mixed 3:1 bilayer of 1-palmitoyl-3-oleoyl-sn-glycero-2-phosphoethanolamine (POPE) and 1-palmitoyl-2-oleoyl-sn-glycero-3-phosphoglycerol (POPG) lipids, a model bacterial-mimicking membrane(24–26). We systematically develop a coarse-grained model for both sequence-isomers of the *β*-peptide within the frame-work of popular MARTINI model. The developed coarse-grained model, in combination with MARTINI membrane(27) and MARTINI polarisable coarse-grained water (28), enabled us to investigate longer time-scale event involving membrane-*β*-peptide interaction. Interestingly, in these ultra-long coarse-grained MD simulations involving multiple copies of GA isomers of AAK and membrane, we observed spontaneous formation of peptide-induced transmembrane pore and water leakage inside the hydrophobic core, leading to disruption of membrane integrity. Analysis based on the tilt angle and end-to-end distance of the pore-forming *β*-peptides suggested that these short-chain biomimetic peptides are able to form trans-membrane pore by stretching themselves across the membrane in a tilted manner. The pore predicted by the coarse-grained simulation was found to be resilient even in an atomically refined model. Finally, the pore forming ability by *β*-peptides was found to be sequence-dependent, with the non GA isomers, at similar P/L ratio, not being able to disrupt the membrane integrity. Taken together, the investigation elucidated a comprehensive mechanism of membrane-disruption by short-chain synthetic biomimetic peptides and identified water-filled pore formation as a key event.

## 2 METHODS AND MATERIALS

### 2.1 Benchmark all-atom simulation

All-atom simulations of membrane-adsorption of *β*-peptide were first carried out as a benchmark for validating the coarse-grained model. Towards this end, a system of single copy of *β*-peptide in a mixed 3:1 POPE/POPG lipid bilayer in water was constructed. For this purpose, the mixed lipid bilayers were first assembled by introducing 292 POPE and 96 POPG lipid molecules (and hence a total of 388 lipid molecules) along with 15722 water molecules using CHARMM-GUI web server(29). The single copy of *β*-peptide was placed in solvent phase. To account for the residual positive charges introduced by the addition of the *β*-peptide (having three *β*-lysine groups) and the charged POPG lipids, in the aqueous media of the membrane, appropriate number ions were introduced so that each of the final systems are rendered charge-neutral. The all-atom system with *β*-peptide and lipid molecules (at a P/L ratio of 1:388) was simulated by 500 nanosecond, the time scale of which was ascertained by visual inspection of peptide getting attached to the membrane surface. Each simulation was repeated five times by varying the velocity seeds. We also had carried out multiple realisations of additional all-atom MD simulations at a higher P/L ratio of 7:128 (7 copies of *β*-peptides with 128-lipid bilayer in presence of 10755 water molecules in a charge-neutral system) for 1 microsecond to get a first-hand knowledge of peptide-membrane interaction in all-atom model at considerably high peptide concentration.

CHARMM36 forcefield parameters (30) were employed to model the all-atom lipid molecules. The CHARMM forcefields, specifically parameterised for these *β*-peptides by Zhu et al., (31, 32) and previously used by Mondal et al. (33), was employed here to model the *β*-peptides. The water molecules are modelled by CHARMM-TIP3P water model(34). CHARMM(30) parameters were also used for the parameters of ions.

All atomistic simulations were performed using Gromacs-2018 software package(35). Each of the simulations was first subjected to an energy-minimisation and subsequently, classical Molecular dynamics were performed at an average pressure of 1 bar and average temperature of 310 K. Each of the components (*β*-peptides, lipids and water molecules) were also coupled separately with the thermostat. The temperature was maintained at the desired value employing Nose Hoover temperature coupling scheme(36, 37) using a coupling constant of 1 ps. A semi-isotropic pressure coupling using Parrinello-Rahman protocol(38) was implemented to maintain the desired pressure of 1 bar. The pressure coupling constants and compressibilities along xy and z directions were 5 ps and 4.5 × 10^5^ bar^−1^ respectively. In all simulations, center of mass motions were removed every 100 steps for *β*-peptides, lipids and water molecules individually. Verlet cutoff(39) schemes were implemented for Lennard-Jones interaction extending upto 1.2 nm while particle mesh Ewald schemes were implemented for treating electrostatic interactions. All bond lengths involving hydrogen atoms of the protein and the ligand benzene were constrained using the LINCS (40) algorithm and water hydrogen bonds were fixed using the SETTLE approach(41). Simulations were performed using the leapfrog integrator with a time step of 2 femto-second and initiated by randomly assigning the velocities of all particles from a Maxwell-Boltzman distribution.

### 2.2 Development of coarse-grained model for *β*-peptide

Due to the difference in the nature of backbone frame-work between *α* and *β* amino acid, the project required us to develop MARTINI-compatible model of GA and non GA isomer of AAK. Towards this end, we followed usual MARTINI protocol for parameterising a new molecule, in this case a 10-residue *β*-peptide. Accordingly, as described in the previous section, we first performed benchmark all-atom simulation of single copy of AAK in bulk water and the simulation trajectories served as the reference.

Figure 2A represents the coarse-grained mapping scheme of GA isomer of AAK. The backbone of *β*-amino acid has one extra carbon atom than natural *α*-amino acid. Accordingly we used a five-to-one bead mapping for backbone of the *β*-amino acid. To mimic the effective size and hydrophobicity of 5:1 mapping (compared to usual MARTINI 4:1 mapping(27)), the nonbonding LJ parameters for backbone were increased to *σ*=0.50 nm (instead of MARTINI backbone value of 0.47 nm) and backbone *E* values were increased to 125 percent of the MARTINI backbone values. Since both GA and non GA sequence-isomers of AAK are known to retain their rigid 14-helical conformations in membrane as well, similar MARTINI backbone nonbonding parameters for *α*-helix, were introduced as the atom types. The bond, angle and dihedral parameters of the backbone of AAK were then iteratively parameterised by performing iterative simulation of coarse-grained AAK in MARTINI polarisable water and comparing the distributions against the all-atom trajectories of the single chain of the *β*-peptide in water, till a best fit was obtained for respective distributions derived from all-atom simulations (see figure S1). The side chain parameters of *βY* and *βK* were directly adopted from analogous MARTINI side-chain parameters of their *α*-amino acid counterparts(42) (compatible with MARTINI polarisable water), as the side chains in both cases are identical. Finally, the cyclic *β*-amino acid ACHC was coarse-grained via 3:1 mapping (usual MARTINI protocol for modelling cyclic functional group) and analogous MARTINI parameters of cyclohexane were adopted for side-chain of ACHC. In order to determine the qualitative accuracy of the coarse-grained model developed for *β*-peptide, we first investigated, if at single peptide level, the coarse-grained model of GA isomer displays qualitatively similar membrane interaction as its atomistic counterpart. Accordingly we compared the adsorption ability of single-copy of GA isomer of AAK on POPE/POPG mixed bilayer in both all-atom and coarse-grained model. We found that the coarse-grained model was able to display spontaneous adsorption process of GA isomer of AAK on the membrane-interface, consistent with all-atom simulation (figure 2B). For a more quantitative validation, we analysed the free energetics of adsorption of the single copy of the *β*-peptide with the membrane interface in both all-atom and coarse-grained simulation. Keeping in the spirit of the current investigation’s theme of using unbiased simulation, we employed a robust Markov state modelling (MSM)(43, 44) approach for computing the free energy of membrane adsorption of single peptide in both representation. PyEMMA (45)(http://pyemma.org) was used to construct and analyse the MSM from all the obtained trajectories. The schematic of the MSM protocol is illustrated in figure 2B. For this purpose, we individually initiated multiple sets of all-atom and coarse-grained MD trajectories of single *β*-peptide in its course of getting adsorbed on the POPE/POPG mixed bilayer.As per the protocol of MSM, the simulation trajectories in both representation (all-atom and coarse-grained), were discretised into 500 micro states, using the minimum distance between the membrane surface and the *β*-peptide as the clustering metric. The discretised MD data were then used to develop a Markov state model. Subsequently, the MSM is lumped into a two-macro-state model by dividing the micro states between ‘adsorbed’ and ‘solvated’ macro states. Finally, the stationary or equilibrium populations of both the macro states are calculated from the MSM.

**Figure 2:**
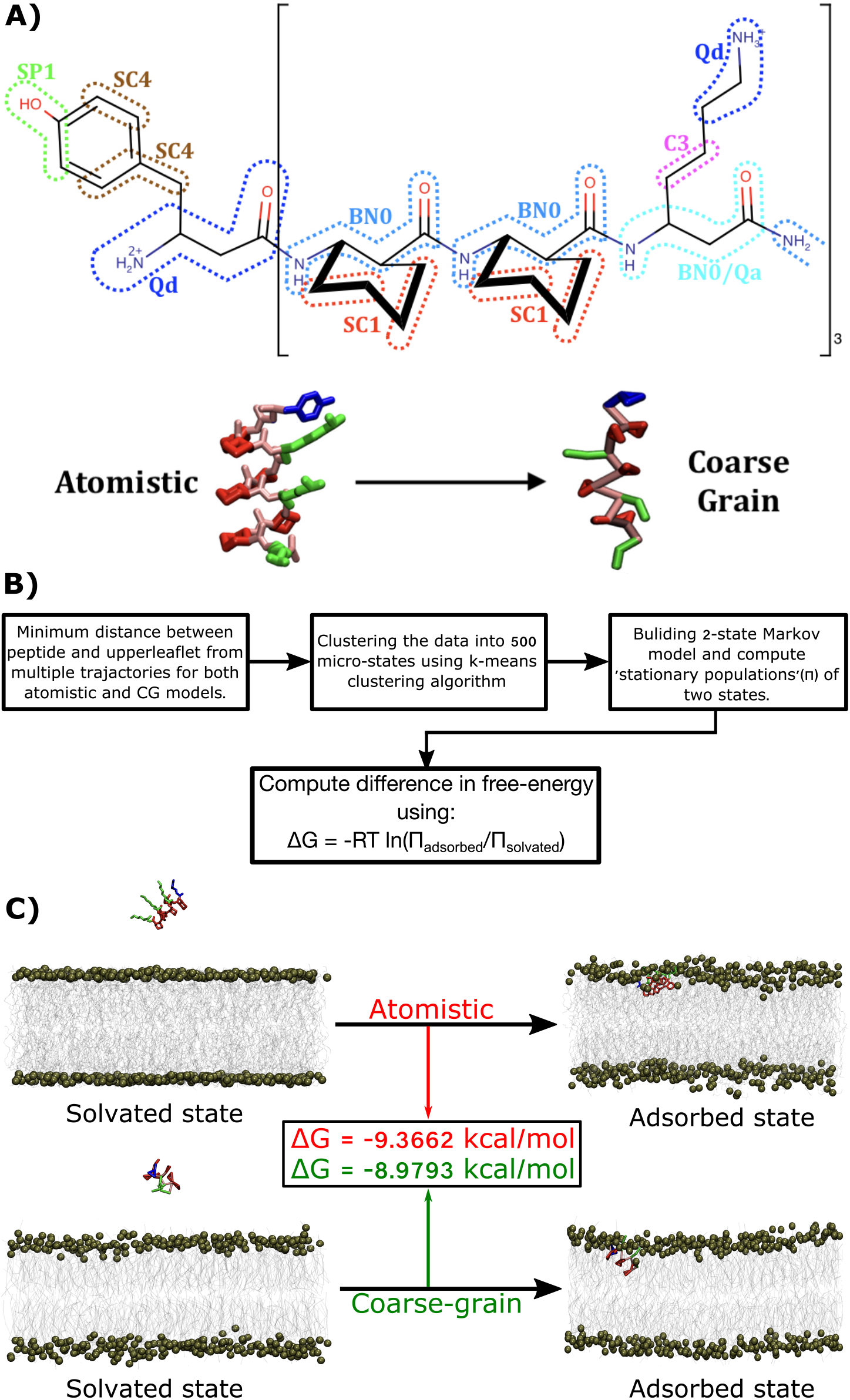
A. Mapping of *β*-peptides to MARTINI based coarse-grained Model. Bottom: Shown also snapshot of all-atom and coarse-grained counterpart. B. Illustration of Markov state model (MSM) scheme for computing the free energy of membrane-adsorption of single copy of *β*-peptide in all-atom and coarse-grained model. C. Comparison of MSM-derived free energy of adsorption at P/L 1:388 between all-atom and coarse-grained simulations.

Adsorption free energies (ΔG) of *β*-peptide to membrane was calculated from the stationary populations of adsorbed and solvated macrostates as obtained from the MSM,

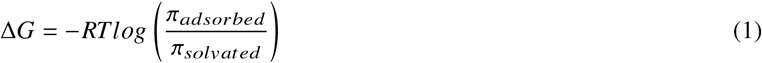

For a systematic comparison, the entire simulation protocol and parameterisation was repeated for non GA sequence isomer as well. We refer to table S1 for system details for simulation using nonGA isomer. See figure S2 for the mapping scheme and optimised distributions.

For reference, we have uploaded topology and parameter files of GA and nonGA isomers of AAK along with an initial configuration of a single AAK in presence of membrane in the Github. The link for downloading the materials : https://github.com/JMLab-tifrh/coarse-grained-models-of-beta-peptides

#### 2.2.1 Long time scale peptide-membrane simulation using coarse-grained model

Table 1 provides a summary of each of the coarse-grained MD simulations carried out with GA isomers of AAK in POPE/POPG lipid bilayers. A total of four separate systems was investigated, using a wide range of P/L ratio which involved 1-15 copies of GA isomer copies. In all CG simulations, we first mapped atomistic protein structure to the coarse-grained model using martinize.py script, appropriately modified for *β*-amino acid. Subsequently, we set up the POPE:POPG (3:1) bilayer along with its solvation by MARTINI polarised water by insane.py (http://www.cgmartini.nl/index.php/downloads/tools/239-insane). In all our systems of interest, simulations were initiated by placing all the *β*-peptide copies in the aqueous phase close to *one* of monolayers of the POPE:POPG (3:1) bilayer, with the empty space filled with martini polarised water ionized with a suitable amount of *Na*^+^ and *Cl*^−^ martini ions to reach an electro-neutral solution. The overlapping molecular contacts were initially removed by energy minimisation using steepest descent algorithm. The asymmetric distribution of the peptides mimics the addition of peptide to a solution containing cells or liposomes, in which peptides adsorb to one monolayer only. Sufficient water layer was introduced on the both side of the bilayer to ensure the *β*-peptide copies do not artificially reach to the other leaflet via periodic boundary condition. The production MD simulations were carried out for 20 *µs*. All production MD simulations were performed at 310.15 K, representing physiological temperature and 1 atmospheric pressure. The average temperature was maintained by velocity rescale temperature coupling scheme with membrane, proteins and water molecules individually coupled to thermal bath with a coupling constant of 1 ps. A semi isotropic pressure coupling approach was employed via Parrinello-Rahman scheme with a pressure coupling constant of 12 ps and isothermal compressibility of 1 bar^−1^. Verlet cutoff scheme was employed for non bonded interaction with a cutoff for Lennard-Jones potential at 1.1 nm. The real space cutoff Coulomb potential was 1.1 nm with relative dielectric constant of 2.5 for polarized water. The long range electrostatic interaction was handled by reaction field, as per so-called MARTINI *New-RF*’ recommendations (46). All MD simulations were performed with the Gromacs 2018 simulation package(35), together with GPU accelerations (46) A time step of 20 femtosecond was used and the neighbour list was updated every 10 steps. The simulation length for each trajectory at higher P/L ratios of 1:39 and 1:22 was 20 *µs*, while at P/L ratio of 1:67, the simulations were extended for 40 *µs*. All simulation times are reported as coarse-grained time-scale. For each P/L ratio, four independent simulation trajectories were spawned. A 5 *µs*-long control simulation with only POPE:POPG lipid bilayers containing 336 lipids (i.e. lacking any peptides) were also performed for comparison purpose.

**Table 1:** Details of the systems for simulation using GA isomer of AAK using MARTINI model. All *β*-peptide copies were initially placed in aqueous phase close to one side of the membrane. Each system was charge-neutralised by adding necessary counter ions.

| System | Number of water | P/L ratio | Simulation Time($\mu s$) × number of replica |
| --- | --- | --- | --- |
| β-peptide-less bilayer | 6538 | 0 | 5 $\mu s \times 2$ |
| 1 GA in 388 lipid | 7761 | 1:388 | 20 $\mu s \times 2$ |
| 5 GA in 336 lipid | 6461 | 1:67 | 40 $\mu s \times 4$ |
| 10 GA in 388 lipid | 10010 | 1:39 | 20 $\mu s \times 4$ |
| 15 GA in 336 lipid | 8324 | 1:22 | 20 $\mu s \times 4$ |

The resilience of the *β*-peptide/membrane complex, as predicted by coarse-grained model, was further assessed by reconstruction of all-atom model via ‘back mapping’ algorithm(47) and subsequently subjecting the refined structure to two independent all-atom 1 *µ*s-long trajectories. Same all-atom simulation protocol, as described earlier (for benchmarking simulation), was implemented.

## 3 RESULTS

### 3.1 An optimised model for membrane-*β*-peptide interaction

To access relatively longer time scale event of interaction between membrane and our biomimetic peptide of interest *β*-peptide, we employed a coarse-grained framework based on MARTINI force field. (27, 42, 48–50) In this work, a mixed composition of phospholipid bilayer of POPE and POPG at 3:1 ratio has been chosen due to its past consideration as a mimic for bacterial membrane(24–26). Additionally, *β*-peptides also have been previously investigated for its membrane activity in experimental assays solely involving model phospholipid bilayers or large unilamelar vesicle(51). The lipid bilayer composition was modelled using MARTINI lipid force field. The choice of coarse-grained model allowed us to use relatively wider lipid bilayer (see table 1) than that typically employed in all-atom simulations. We employed polarisable MARTINI water model (28) for coarse-grained MARTINI force field for simulating the aqueous phase of the lipid bilayers. The choice was due to its superior ability to reproduce the water dielectric property, orientational polarizibilities and membrane interfacial electrostatic property more faithfully than the regular MARTINI water model. Accordingly, the Martini lipid and protein force fields compatible with MARTINI polarizable water were chosen in the current work. As detailed later, we would find that the usage of polarisable MARTINI water(28) (in contrast to regular MARTINI coarse-grained water(27)) was instrumental in observing peptide induced trans-membrane pore in long time scale simulations. Due to the difference in the nature of backbone frame-work between naturally occurring *α*-amino acid and synthetically designed *β*-amino acid, the project prompted us to design coarse-grained model of GA and non GA isomer of AAK. As detailed in the method section, we developed a MARTINI-compatible coarse-grained model for the 10-residue *β*-peptide AAK, by iterative optimisation against all-atom simulation trajectory. Figure 2A provides the coarse-grained mapping scheme employed in the current work for GA isomer of the *β*-peptide.

Both in vitro and in vivo investigations into the anti-fungal and antibacterial abilities of sequence-isomers of *β*-peptide of our interest, AAK, have shown that it kills pathogen cells at a much lower concentration than many naturally occurring antimicrobial peptides. Towards this end, we first explored how the membrane/*β*-peptide interaction behaviour at a *single peptide level*, simulated using coarse-grained frame-work, would compare against those performed using an atomically detailed force field. We individually simulated the interaction of single copy of GA isomer of AAK, initially present in aqueous media, with a 3:1 POPE/POPG phospholipid bilayer, in an all-atom and coarse-grained representation. We find that, in both all-atom and coarse-grained simulations, single copy of AAK, starting from aqueous media, gradually approaches the hydrophilic interface of the phospholipid bilayer and eventually gets partitioned in the inner membrane interface. Representative snapshots in figure 2C provide a pictorial view of the solvated and eventual membrane-adsorbed state of GA isomer of AAK at single peptide level, obtained from all-atom and coarse-grained model. The comparison indicates qualitatively a very similar membrane-adsorbed state obtained in both types of simulations at single peptide level. For a quantitative comparison of the quality of coarse-grained model, as against atomistic model, we computed the free energy of *β*-peptide’s membrane-adsorption in both models. As described in method section, we adapted a framework of MSM (see figure 2B for details of protocol), based on multiple unbiased simulation trajectories, to compute the relative stationary or equilibrium populations of two macro states of *β*-peptides, namely ‘solvated’ and ‘membrane-adsorbed’state. The stationary populations of the ‘solvated’ (*π*_*solvated*_) and ‘adsorbed’ (*π*_*adsorbed*_) states, as derived from MSM underlying the all-atom model, were found to be 0.0142 % and 99.9858 % respectively. On the other hand, MSM built using coarse-grained simulation predicted a value of *π*_*solvated*_= 0.0273 % and *π*_*adsorbed*_= 99.9727 %. Accordingly, the free energy of adsorption, as computed by equation 1, turned out to be -9.37 kcal/mol in atomistic simulation, as compared to -8.97 kcal/mol in coarse-grained model, indicating reasonable agreements. Together, these observations validate that the coarse-grained model is able to capture the basic membrane-adsorption feature of amphiphilic *β*-peptide very well. Moreover, at high peptide concentration, we find that at short-time scale, the present coarse-grained model reproduces very similar early-stage behaviour of membrane/*β*-peptide interaction, as found in 1 *µs*-long all-atom simulation of membrane interactions with multiple copies of *β*-peptides: In both cases (see first two rows of figure 3 for coarse-grained simulation and figure S3 for all-atom simulation) the *β*-peptides undergo spontaneous self-aggregation and membrane-adsorption within a 1 *µs* long simulation. These initial benchmark assessments of the coarse-grained model provided early promises for employing them in long time scale investigations.

**Figure 3:**
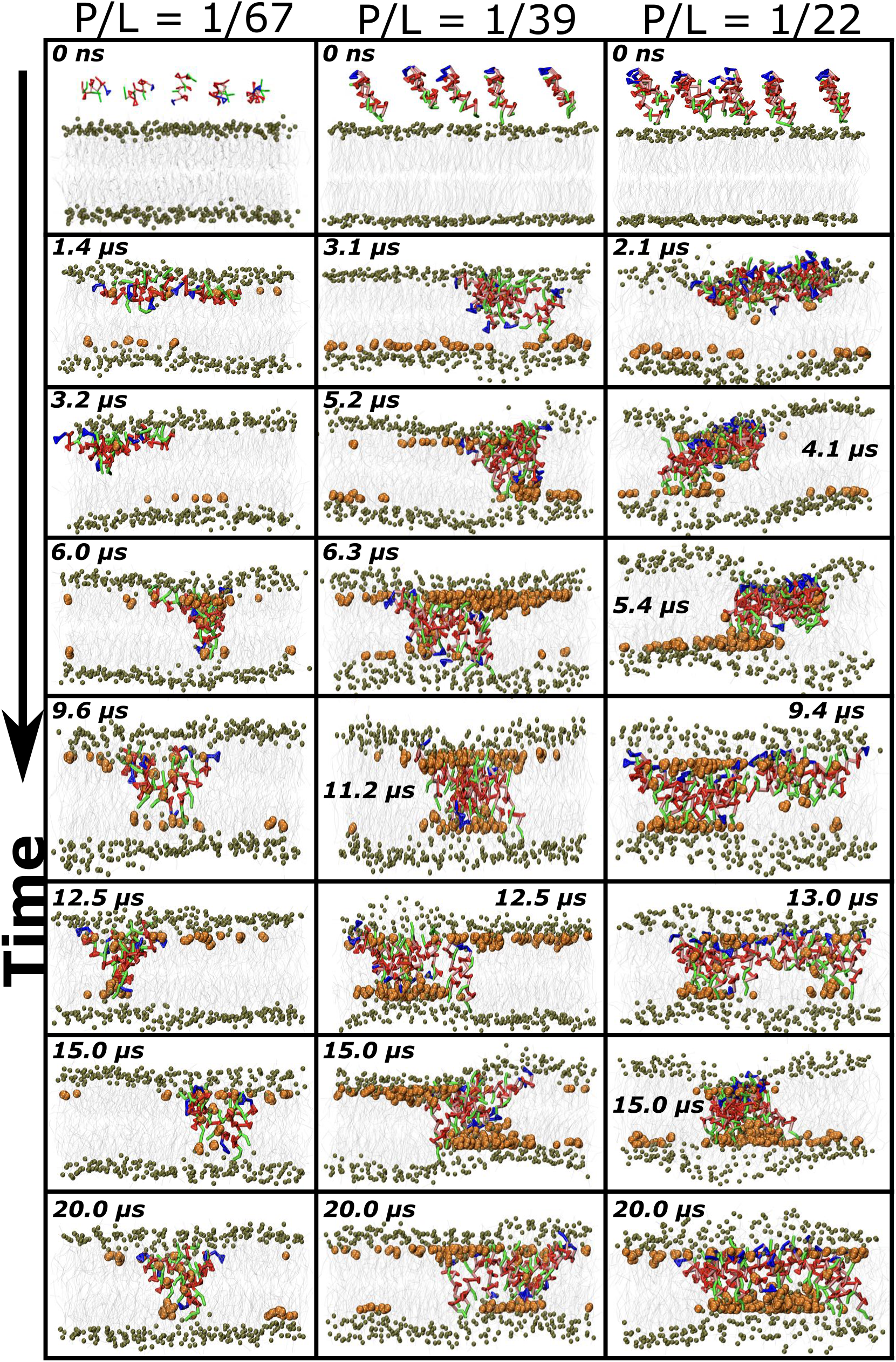
Time profile of account of *β*-peptides/membrane interaction at diverse P/L ratios. As a general trend, the *β*-peptides, after initiating from aqueous media, gradually assembles and partitions into the membrane interface. This is followed by transmembrane insertion of a number of *β*-peptide copies inside the membrane. The *β*-peptides eventually span across both the bilayer. Color code: tan : Phosphate groups of lipids, orange : Polarized water, blue :: Tyrosin, green : Lysin, red : ACHC residues.

### 3.2 Simulations capture spontaneous transmembrane pore formation by short-chain *β*-peptides

Encouraged by the observation of efficient membrane-adsorption GA isomer of AAK towards membrane interaction manifested at single *β*-peptide level in a coarse-grained model, we explored the effect of high concentrations of GA isomer on the membrane. As enumerated in table 1, in our coarse-grained simulation, we systematically varied the P/L ratios within a wide range of 1:388 to 1:22 to investigate the interaction of increasing concentration of *β*-peptides with the membrane. At a very low P/L ratio of 1:388, which involved interaction of single copy of AAK with the membrane, as described earlier, the *β*-peptide approached the membrane interface of the proximal leaflet and remained partitioned across the membrane interface. However, intriguingly, at a threshold P/L ratio of 1:67 (containing 5 peptides and 336 lipid molecules), coarse-grained simulations at longer coarse-graining time scale up to 40 *µs* displayed that (see movie S1 and left column of figure 3 for a visual account of process) few copies of GA isomer of AAK, initially mutually well-separated in bulk water gets self-assembled and adsorbed on the membrane surface. The aggregates of *β*-peptides, upon being docked on the surface, act as a cage and displace a number of lipid head groups (see figure S4 A). The lipid displacement provides the required opening for favorable hydrophobic interaction between ACHC residues and the lipid tails and as a result the *β*-peptide copies got past the polar head group of the proximal leaflet of the membrane interface. The aggregation and organisation process of the *β*-peptides continue past the event of membrane-adsorption. Eventually, the peptide-aggregate, upon attachment to membrane interface, snorkelled past the membrane interface of proximal leaflet and got embedded in a transmembrane orientation and spanned across proximal and distal leaflets of the membrane. As seen in movie S1 and in left column of figure 3, the defects created due to peptide insertions across the membrane interface led to a number of water molecules getting leaked inside the membrane interiors, thereby giving rise to a transmembrane pore-like structure for maximising favourable contacts between lipid tails and the hydrophobic *β*-amino acid residues.

### 3.3 A cooperative pore formation process

We systematically increased the P/L ratios to 1:39 and 1:22 by introducing additional copies of the GA isomers in the same side of water phase and performed series of individual MD simulations using coarse-grained model. Movie S2 and S3 demonstrate the event of the *β*-peptide/membrane interaction at the P/L ratios of 1:39 and 1:22. Figure 3 provides a visual account of membrane-peptide interaction at different time points of multi *µs* long trajectories at P/L ratios of 1:67, 1:39 and 1:22. In all these three P/L ratios, the trend of pore formation is generic, except that the event gets accelerated at higher P/L ratio. At P/L ratio of 1:22, the formation of two distinct trans-membrane pores is also seen at an intermediate time point, which eventually coalesce into a single large pore. The pore formation event was found to be reproducible across multiple trajectories (2 out of 4 trajectories for P/L=1: 67 and 3 out of 4 trajectories for P/L=1:39 and 1:22 displayed stable pore formation. Each trajectory was 20-40 *µs* long.) As shown in figure S5, at lower than five peptide copy numbers and at a P/L ratio lower than 1:67, the peptides would only settle below the surface of bilayer causing a partial disruption. This indicates that pre-association of the peptides may increase pore propensity as shown experimentally in case of tetravalent magainin peptides(52). Once the formation of transmembrane pore is completed, the pore remains stable for the rest of the period of simulation. On the basis of the observation that pore formation is facilitated by self-aggregation of *β*-peptides and a threshold P/L ratio is required for transmembrane pore formation, we conclude that this is a cooperative process. This is qualitatively consistent with past free energy based simulation result, albeit focussed on naturally occurring antimicrobial peptide melittin, that increasing the number of peptide copies reduces the barrier to pore formation(53).

### 3.4 Quantifying and characterising the pore formation process

To characterise the extent of *β*-peptide insertion inside the membrane at a given P/L ratio, we compute the minimum distance (*d*_*min*_) between individual peptide copies and the phosphate group of the distal leaflet of the membrane (see schematic in figure 4A). As shown in the figure 4A, for single peptide level, *d*_*min*_ remains stable at a value of 2.5 nm, which corresponds to the peptide remaining adsorbed in the membrane-interface of the proximal leaflet at a P/L of 1:388. However, at higher P/L ratios of 1:67, 1:39 and 1:22, we observe a sharp collective drop in the value of *d*_*min*_ from 2.5 nm to 0.5 nm for a number of peptide copies at a time scale of 5-10 *µs* in coarse-grained simulation, indicating the onset of translocation of a cluster of peptides inside the membrane interior and initiation of eventual transmembrane pore formation (left column of Figure 4). Interestingly for P/L ratio of 1:22, we observe two such events in which *d*_*min*_ decreased for two independent peptide clusters, indicating the onsets of two distinct pore formations as confirmed by Movie S3 and of top panel of Figure 4A. Once the transmembrane pore formation is completed, the pore remains stable for the rest of the duration of 20 *µs* (40 *µs* for P/L ratio of 1:67) simulation.

**Figure 4:**
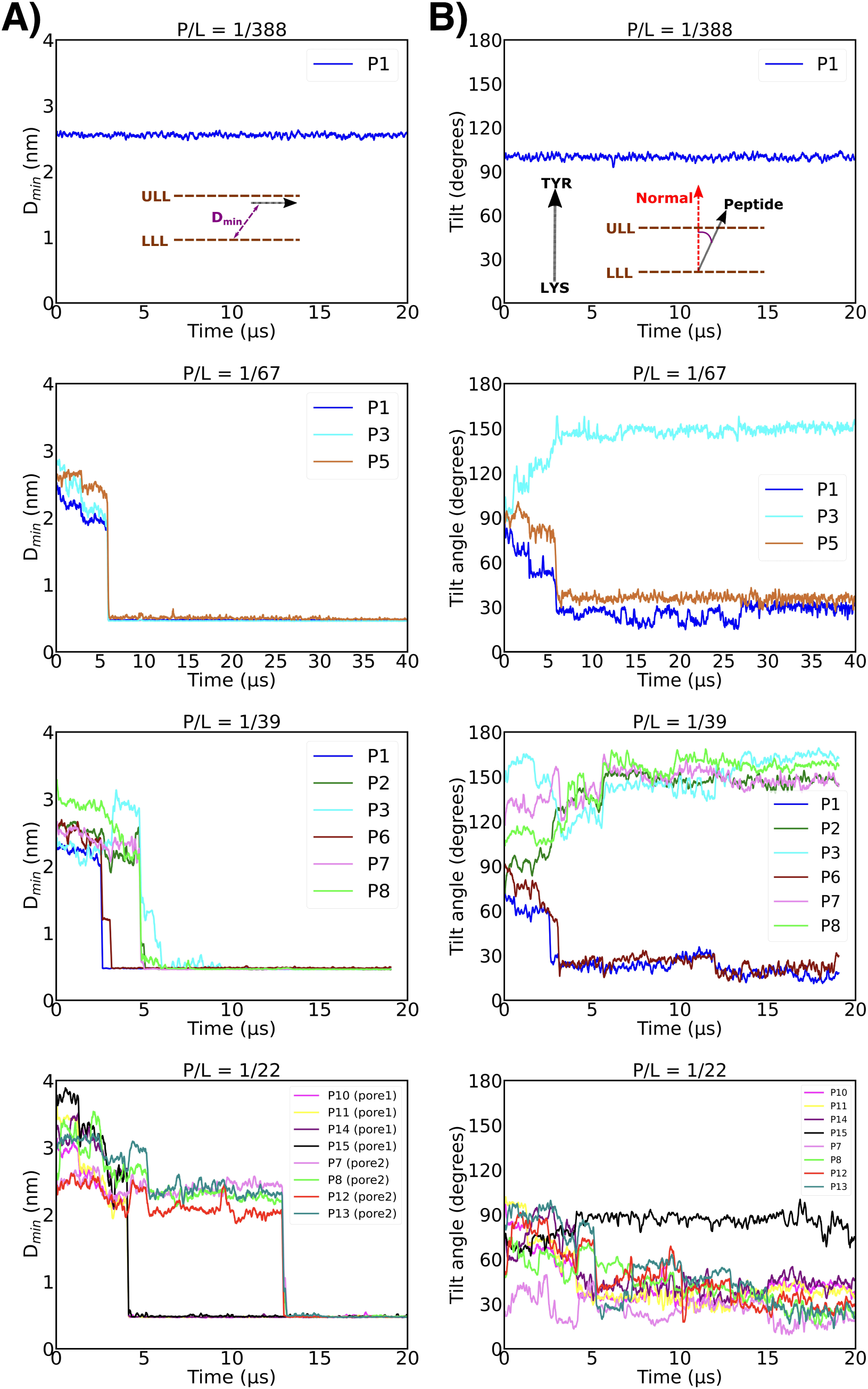
A. Comparison of time profile of extent of peptide insertions. Here, minimum distance between the phosphate group of distal leaflet and the peptide (*d_min_*) has been considered as the metric. The sudden drop in the value of *d_min_* signifies the onset of peptide translocation and pore formation. Only the time profile of pore-forming peptide copies (annotated as ‘P1, P2…’) has been shown. B. Comparison of time profile of tilt angle of the peptides with the membrane interface at diverse P/L ratios.

Figures 4 B characterise the tilt angle between the *β*-peptide copy and the membrane surface normal. Based on this analysis, a tilt-angle of 90 degree would imply that the peptide is horizontally oriented with the membrane interface, which is the case for single peptide at P/L of 1:388. However, with increasing P/L ratio, we found that the *β*-peptides which formed the core of the pore, eventually transitioned from a tilt angle of 90 degree to a tilt angle closer to 30 degree or 150 degree (suggestive of reversal of peptide’s orientation during membrane translocation), indicating that these pore-forming peptides have attained a transmembrane orientation. However, the tilt angle is not perfectly zero, suggesting a tilted orientation subtended by these pore-forming *β*-peptides with the membrane interface. Together, the analysis of time profile of tilt angle and *d*_*min*_ of the peptides indicate that the formation of transmembrane pore involves collective translocation of *β*-peptides at a multi-microsecond long time scale, in which this peptide eventually makes a tilted orientation with the membrane interface. The timescale required for the formation of pore containing minimum of five peptides (at a P/L ratio of 1:67) was around 8-9 *µs* in coarse-grained unit. As the P/L ratio and number of *β*-peptide copies increased to 1:39 and 1:22, the pore formation process and reorientation to transmembrane pose get expedited (2.5-5 *µs*). The reorientation of peptide from a surface to a transmembrane conformation is generally considered an important step in its pore-forming ability(54, 55) and is believed to be associated with a considerable free energy barrier(56). The observation of expedited transmembrane reorientation of *β*-peptides beyond a threshold P/L ratio qualitatively coincides with the previous report of reduced free energy barrier of reorientation of melittin with increasing P/L ratio(56). Overall, similar analysis based on distance and tilt-angle also characterises transmembrane insertion of peptide clusters across multiple trajectories (see Figure S6 and S7 for P/L ratio of 1:39 and 1:22)

Figure 5A compares the peptide density profiles along the surface normal for all P/L ratios investigated here. These density profiles were averaged over last 3 *µs* duration of the simulation (see snapshot in figure 5B). Consistent with previous section’s description, with increasing P/L ratio, we find that the *β*-peptide density past the phosphate group gradually increases inside the hydrophobic core of the membrane and eventually the tail of the density profile stretches upto distal leaflet of the membrane at high P/L ratio, implying that the peptides span both the leaflets and form a transmembrane channel. *β*-tyrosine residue mostly remains localised close to the membrane interface while the hydrophobic ACHC groups span the core of the transmembrane pore. The density of the phosphate groups (figure 5C) also increases inside the hydrophobic core of the membrane with increasing P/L. The sharp phosphate density profile in neat membrane gets gradually smeared with increasing P/L ratio. Most importantly, as is observed in the water density profile (see figure 5C), the peptide-membrane interaction leads to considerable increase in water density inside the hydrophobic core of the membrane with increasing P/L ratio. The representative snapshots at high P/L ratios of 1:39 and 1:22 especially depict (figure 5B) the presence of a distinct layer of water near the inner part of the both the leaflets. This suggests that these transmembrane pores are wet in nature and we believe that these interfacial waters help maintain the transmembrane pore via favourable interactions with the polar terminal residues of AAK.

**Figure 5:**
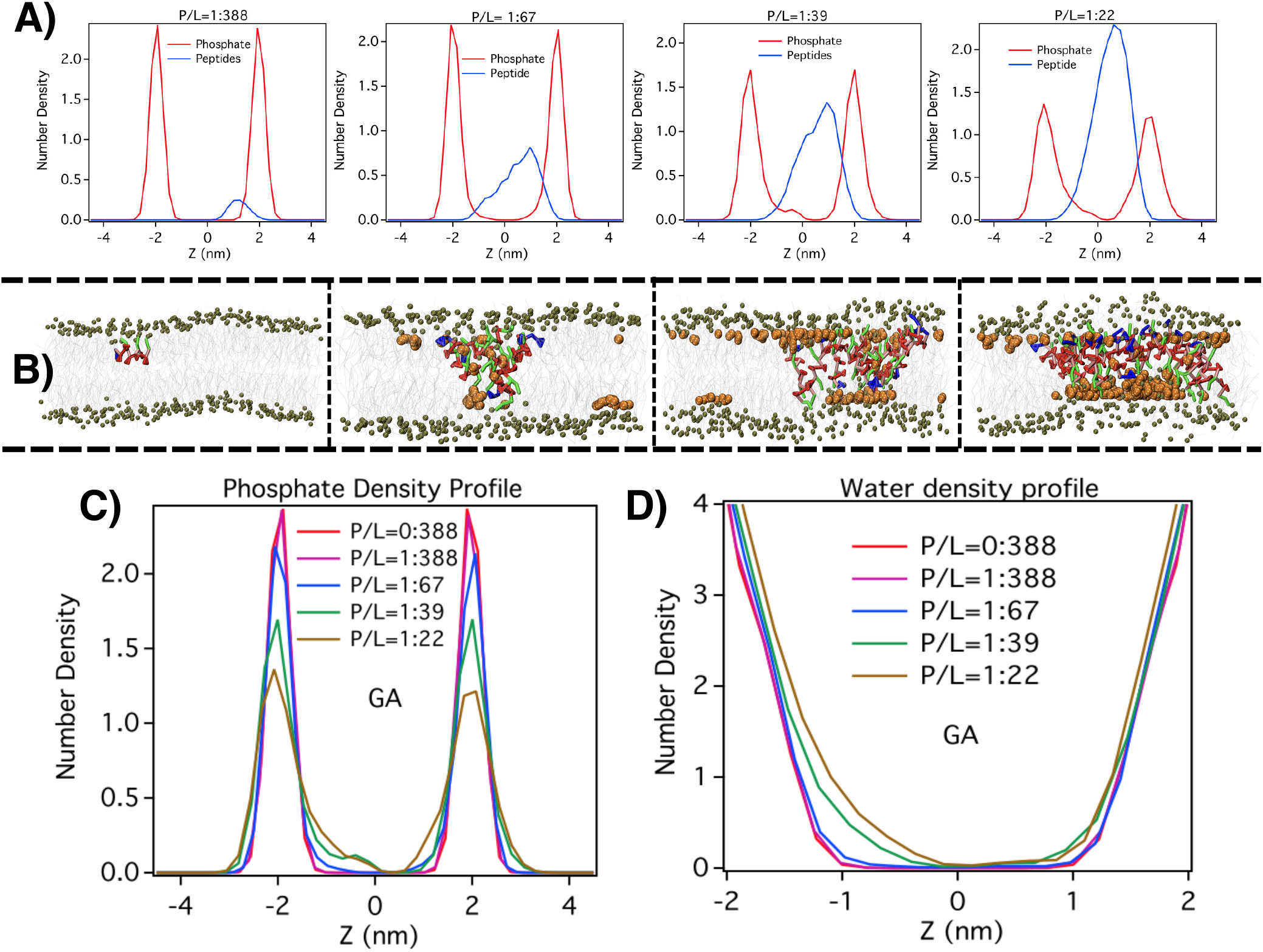
A. Density profile of GA peptides across membrane interface at multiple P/L ratios. B. shown are the representative snapshots of equilibrium conformation in all P/L ratio. C-D: Density profile of phosphate and waters at multiple P/L ratios.

Figure 6A quantify the extent of water leakage till the membrane centre at two high P/L ratios of 1:39 and 1:22 (in which the pore formation was most prominent). We find that a considerable number of water molecules get past the membrane leaflets and the number of leaked water molecules systematically increases as the event of peptide-induced pore formation proceeds. The trend of water leakage is also higher at larger P/L ratio. The simulation trajectories were quantitatively analysed for exploring the possibility of occurrences of trans-membrane water permeation events, in which a water molecule, after starting from one membrane leaflet, would be tracked for its completing the full passage to the other leaflet. Figure 6B compares the cumulative number of such water permeation events though the membrane at two P/L ratio. The trend reveals that there are in fact, multiple occurrences of water permeation event in which water molecules traverse from one leaflet to other leaflets and the extent of water permeation increases with increasing P/L ratio. Figure 6C illustrates a time profile of two representative water permeation events across the two leaflets. An analysis of water permeation trajectory indicated that the duration of water permeation event can be diverse. The distribution of duration of water permeation events at two P/L ratios (figure 6D-E) shows that while the majority of the trans-membrane water passage completes in 10 ns, there are numerous occurrences of considerably slow permeation events, as represented by long tail of the distributions.

**Figure 6:**
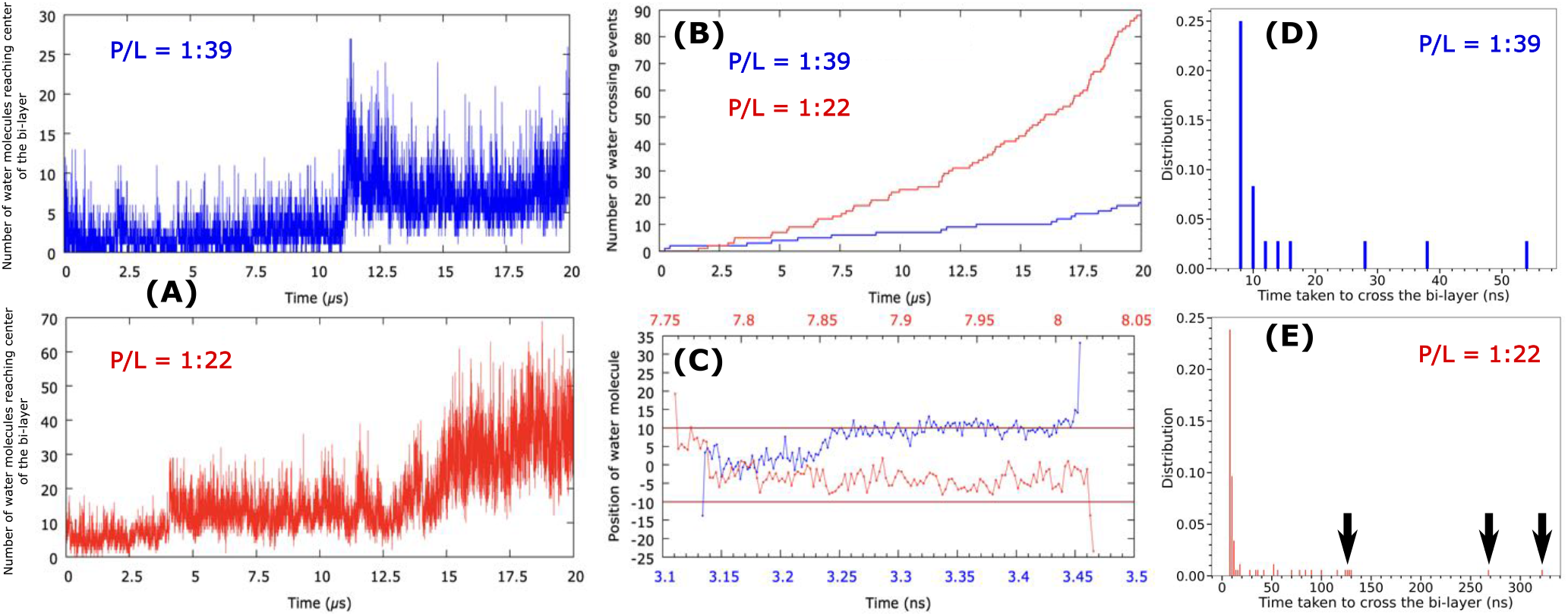
A. Time profile of number of water leaked upto the membrane centre at two higher P/L ratios (1:39 and 1:22).B. Cumulative number of water permeation events during the simulation period at two P/L ratios. Only those water molecules which made a full pass from one leaflet to the other leaflet was counted. C. Trajectory of two representative water permeation events across the leaflet, D-E. The distribution of permeation-time at two P/L ratios.

There are diverse proposals of variants of trans-membrane pores including barrel-stave, toroidal and carpet types (3, 4, 18, 19). The detailed characterisation of the pore showed that *β*-peptides induce local defects in the bilayer and subsequently the head groups of the lipids line the pore together along with the peptides. The visual inspection of the pore also revealed that the peptides are unevenly distributed i.e. higher concentration of peptides were present in the upper leaflet in comparison to the ones in lower leaflet. The tilt-angle analysis of the pore-forming peptides (discussed in the previous paragraph) indicates that the *β*-peptide aggregate gets inserted almost vertically into the membrane, almost baring the characteristics of a barrel stave pore. However, the pore is not perfectly cylindrical, rather it subtends a non-zero angle with the membrane normal. Additionally, the snapshots of the pore also shows minor line-up of the displaced lipid head groups. However, the lining of lipid head-groups is not as prominent as what would be observed in an ideal toroidal pore.

Finally, we stress that the choice of polarizable MARTINI coarse-grained water(28) for modelling solvent was instrumental in observation of the pore in the current simulations. The usage of regular Martini coarse-grained LJ water model only resulted in the peptide adsorption on the membrane leaflet and did not lead to any pore formation. (see figure S8) This suggests the importance of appropriate representation of the membrane interfacial electrostatics via choice of better water model.

We find that the impact of pore formation and water leakage on the overall membrane integrity is mostly local. This is demonstrated in the two-dimensional bilayer thickness profiles (figure 7) at diverse P/L ratios. Within P/L ratio of 1:388 to 1:67, the bilayer thickness profile remains uniform, similar to that in peptide-less membrane. However, at higher P/L ratio of 1:39, we find localised non-uniformity in the membrane-thickness profile, which eventually grows at P/L ratio of 1:22. As an indication of disrupted membrane integrity induced by GA isomer of *β*-peptide, the deuterium order parameters of the POPG lipid tails systematically decrease with increasing P/L ratio. Interestingly, the order parameter of POPE lipid does not change in presence of peptides, indicating specific interaction between POPG lipids and *β*-peptide.

**Figure 7:**
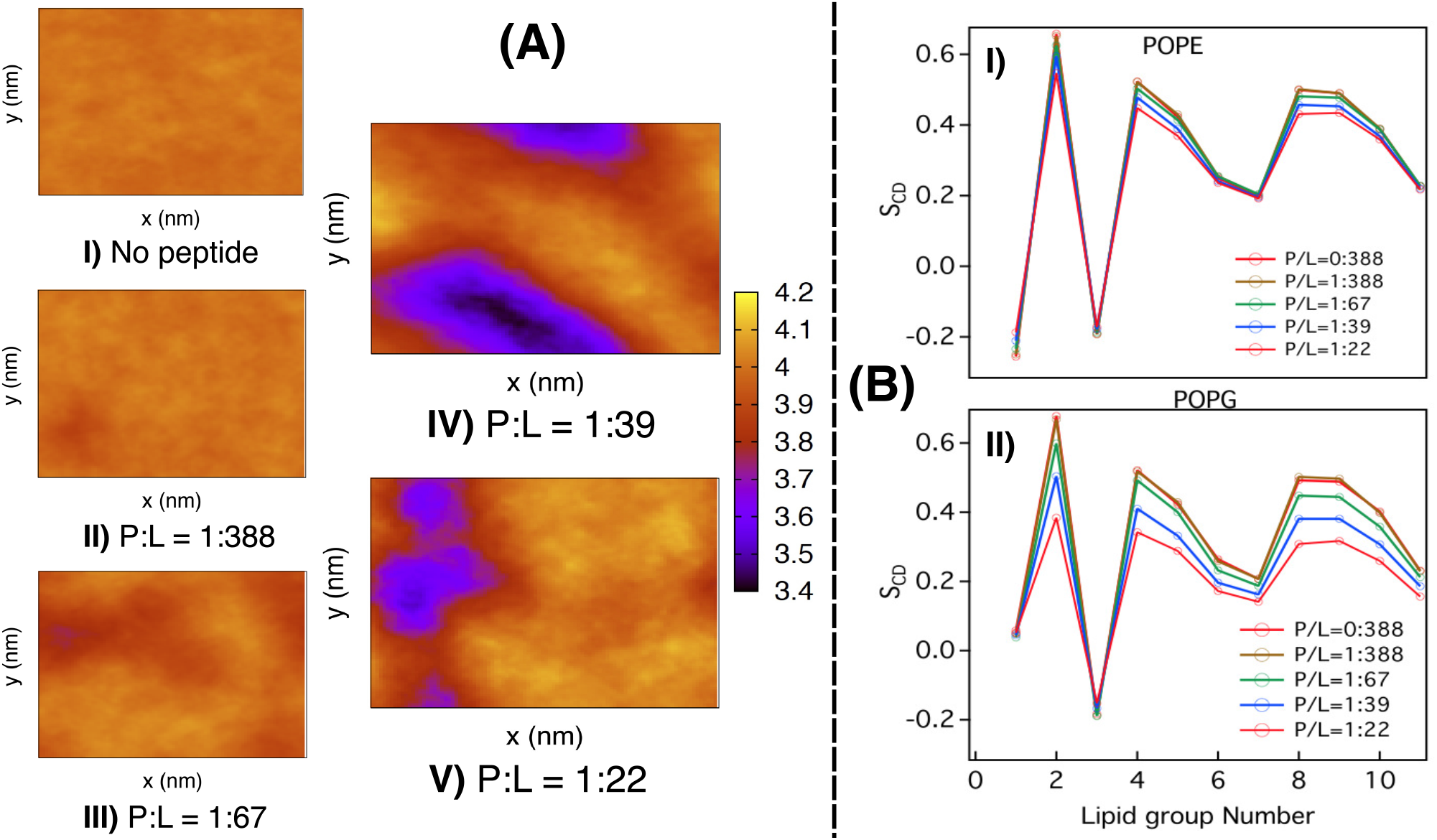
Comparison of A. Bilayer thickness profile B. Lipid order parameter of the POPE and POPG lipids across multiple P/L ratios of GA isomer

### 3.5 Reconciling short-chain architecture of *β*-peptides and its transmembrane pore formation ability

A distinct feature of the *β*-peptide of current interest i.e. GA isomer of AAK is their relatively shorter architecture (only 10-residue) than typical natural antimicrobial peptides such as magainin (23 residues), melittin (27 residues) or LL37 (37 residues). A pertinent question is: how are these short-length *β*-peptides able to form membrane-spanning pore in the bilayer? Towards this end, we analyse the end-to-end distance of the *β*-peptide in different media. Figure 8 A compares the distribution of end-to-end distance of this specific *β*-peptide. We find that in bulk water, the average end-to-end distance is approximately 1.5 nm, which remains almost similar in the case where single peptide remains adsorbed in the membrane/water interface (i.e. at P/L = 1:388). However, at higher P/L ratios, we note that the end-to-end distance of individual transmembrane peptides, which are constituent of the pore, increases to approximately 2.5 nm over time.(figure 8 A) A careful inspection of time profile of the end-to-end distance in figure S9 suggests that the onset of increase in the pore-forming *β*-peptides’ end-to-end distance coincides with the onset of transmembrane pore formation of the *β*-peptides inside the membrane. The extended chain length of these peptides inside the membrane interior enable them to span the membrane and form trans-membrane pore.

**Figure 8:**
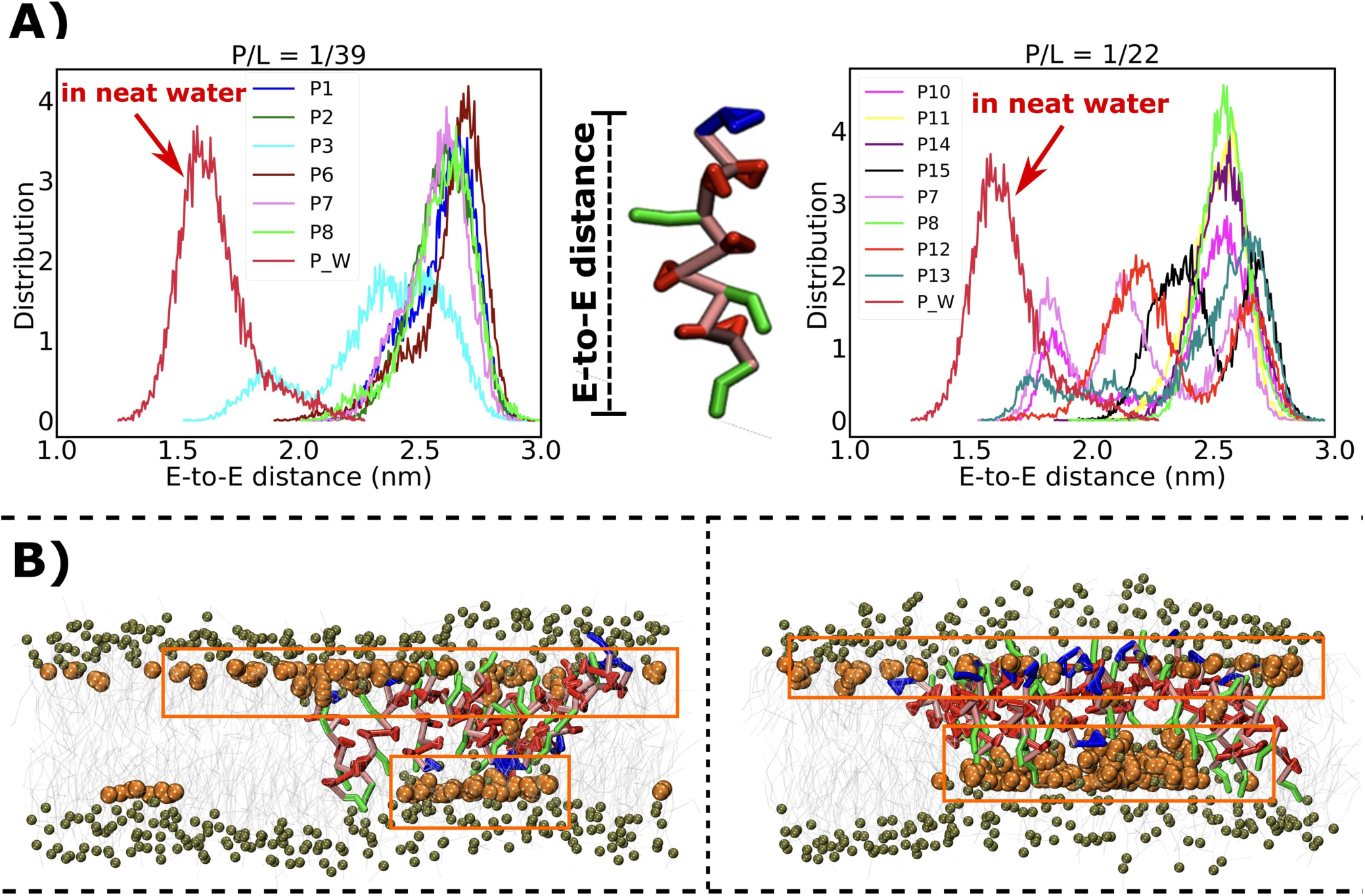
A. Comparison of the probability distribution of GA isomer’s end-to-end distance between that in neat water and that inside membrane at multiple P/L ratio B. The representative snapshot of pore showing water-layer anchoring the transmembrane pores.

A plausible rationale for these observed extension of these pore-forming peptides is that, once inside the hydrophobic core of the membrane, the polar N-terminal residue *β*-tyrosine or hydrophilic C-terminal residue *β*-lysine residue of these *β*-peptides aim to reach out to the polar head groups of the distal leaflet for favourable interaction (see figure S4 B), which leads to a possible elongation of these peptides. Although, the transmembrane *β*-peptides are stretched inside the bilayer, it is still shorter than average membrane thickness of the neat POPE/POPG bilayer. A visual inspection of the transmembrane pore and analysis of bilayer thickness profiles at high P/L ratios of 1:39 and 1:22 (Figure 8 B) shows that the bilayer becomes non-uniform in presence of these *β*-peptides and undergoes localised thinning, which helps these extended pore-forming *β*-peptides to span the leaflet. Finally, as discussed earlier, the presence of layer of interfacial water acts (figure 8 B) as additional cushion to support the transmembrane pore.

### 3.6 A refined model of the pore

The current prediction of formation of trans-membrane pore by these short chain peptides is albeit made in a coarse-grained model, which might be argued for lack of chemical details. In order to assess the resilience of the predicted pore, we reverse-mapped a representative conformation of coarse-grained morphology of the water-filled pore into its full all-atom representation via ‘backmap algorithm’ (47) and subsequently subjected it to relaxation via MD simulation. Figure 9 depicts the refined all-atom representation of peptide in complex with membrane, as obtained after equilibration. We find that the structure of the pore is retained in the reconstructed model. An expanded view of the figure of the refined model confirms the presence of large density of water and displaced phosphate head groups inside the pore. We find that the extended conformation of the *β*-peptides retains its stability in the membrane interior, when subjected to microsecond-long all-atom MD simulation as well, which further validates that the extended conformations of the *β*-peptides inside the membrane core, as predicted by coarse-grained simulation is a realistic possibility.

**Figure 9:**
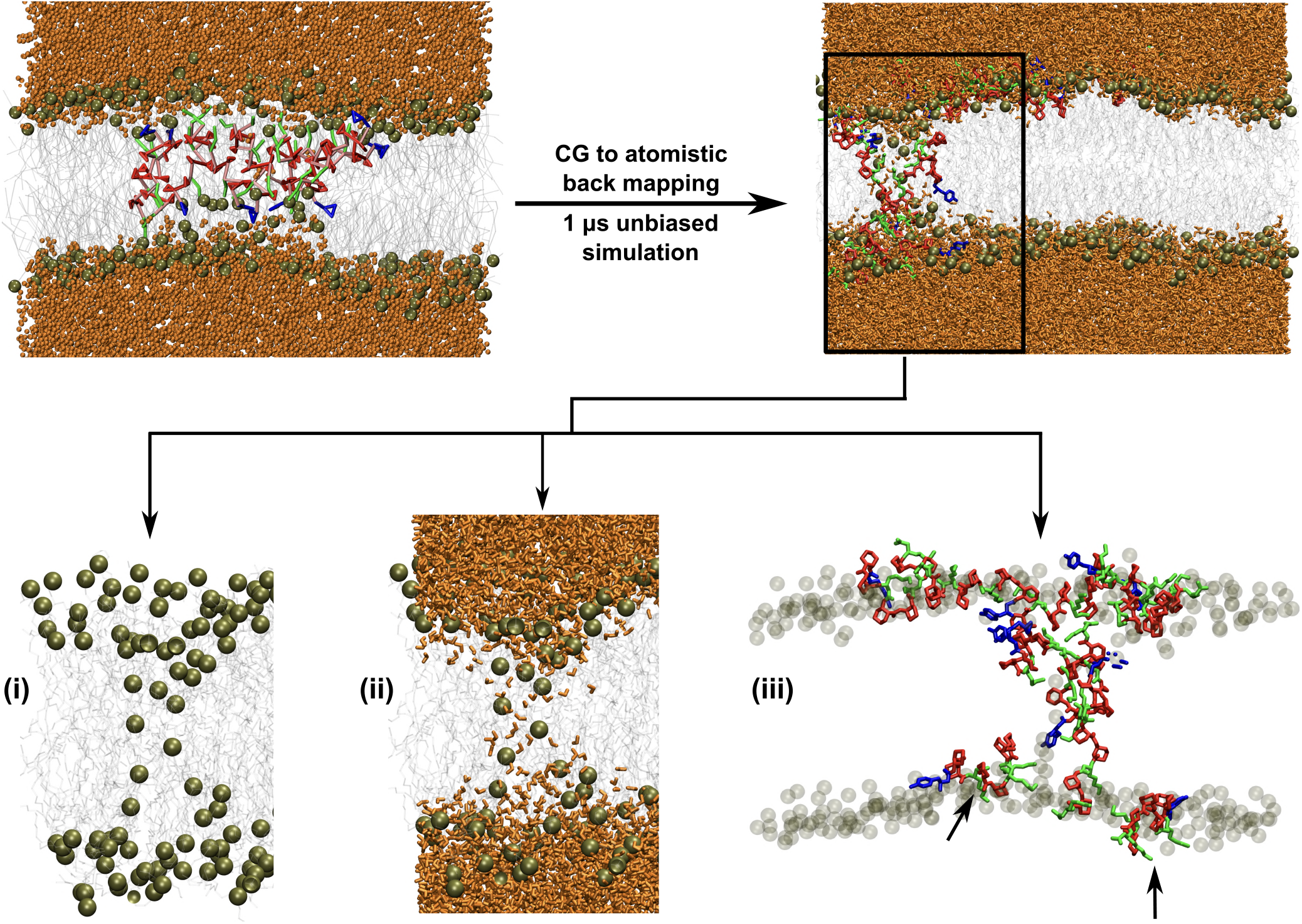
All-atom refined model of the peptide/membrane complex formed at P/L of 1:39, as obtained via reverse-mapping the coarse-grained structure and subsequent micro-second long MD simulation. The presence of water and displaced phosphates in the membrane interior is evident in the expanded image (see inset i and ii). Also shown are the expanded image of the eventual conformation of the *β*-peptides in the all-atom representation. The extended conformation of the peptides are apparent in the all-atom simulation.

### 3.7 The pore formation ability of *β*-peptide is sequence-selective

We investigated whether the pore forming propensity of *β*-peptides is sequence dependent. Accordingly, we also simulated the interaction of non GA isomers of AAK with the membrane at a wide range of P/L ratio, similar to GA isomer. The peptide density profiles of non GA isomers are compared in figure 10A across P/L ratios ranging between 1:388 to 1:22. We find that, unlike GA isomer, the peptide density profile does not span across both the leaflets in non GA isomer even in the highest P/L ratio of 1:22. Instead, as shown by the representative snapshot in figure 10B, we see that the *β*-peptide copies in non GA isomer self-assemble into small isolated oligomeric cluster and remain adsorbed on the membrane interface, without showing any tendency of translocation inside the membrane interior. This is also reflected in unchanged phosphate density profiles and water density profiles in the membranes across entire range of P/L ratio involving non GA isomers (figure 10 C-D). Finally, relatively unchanged bilayer thickness profiles (figure S10 A) and deuterium order parameters (figure S10 B) across P/L ratios in case of non GA isomer relative to GA isomer clearly suggests that non GA isomer of AAK would not be able to disrupt the membrane integrity unlike GA isomer at similar P/L ratio. These results clearly suggest that the presence of global segregation of hydrophilic and hydrophobic residues in a *β*-peptide, as in GA isomer, is key to its ability to form water-lined transmembrane pore and to disrupt membrane integrity.

**Figure 10:**
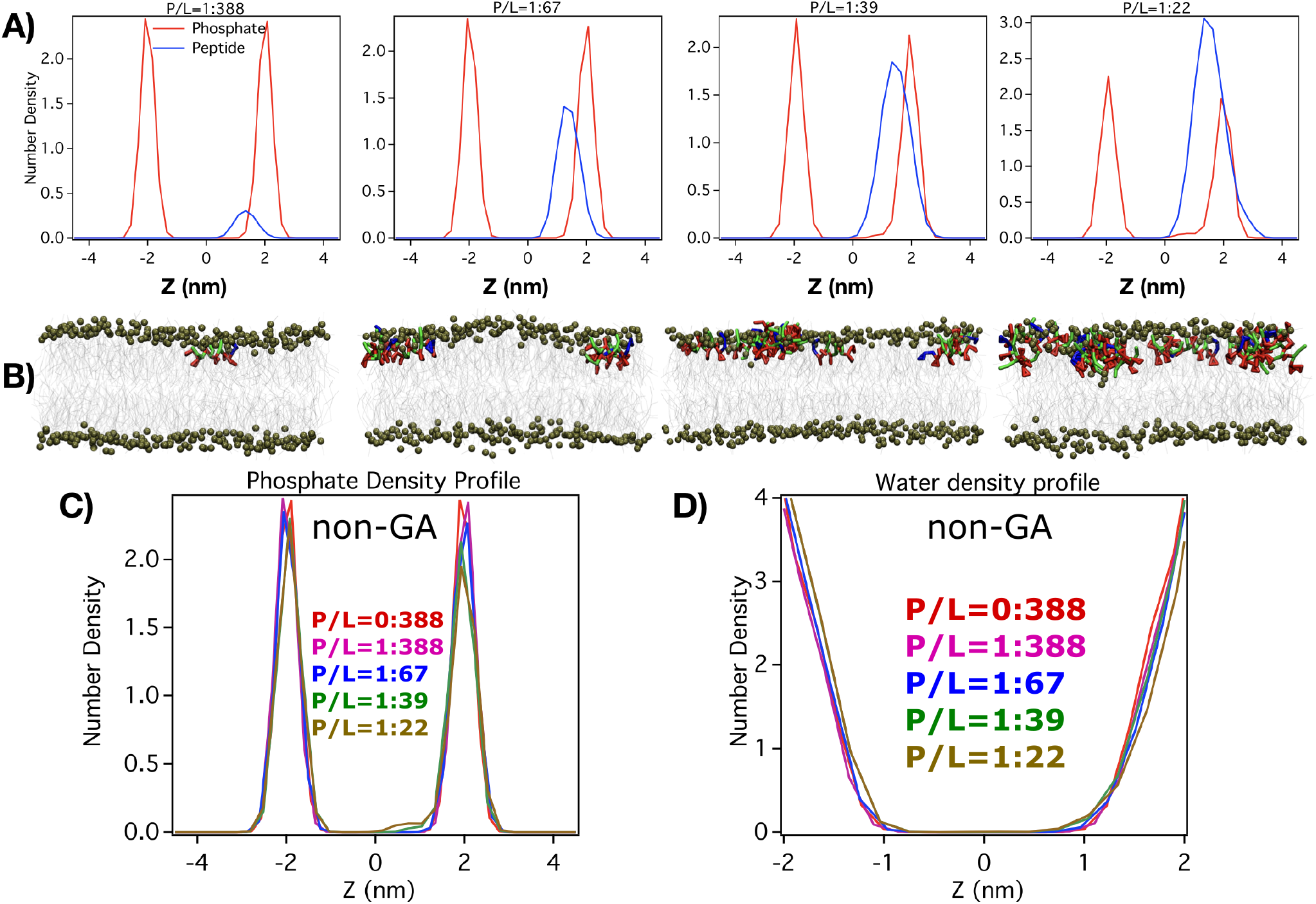
A. Density profile of non-GA isomers across membrane interface at multiple P/L ratios. B. Representative snapshots of non-GA isomers interacting with membrane at all P/L ratios C. Comparison of phosphate density profiles in presence of non-GA isomers at diverse P/L ratios. D. Comparison of water density profiles in presence of non-GA isomers at diverse P/L ratios.

## 4 DISCUSSION

Unravelling the mechanistic action of antimicrobial agents at an atomistic resolution has always been a key desirable from both experimental and computational perspectives. Specifically, it remains more intriguing to decipher the membrane-disrupting mechanism for amphiphilic biomimetic peptides which manifest strong antimicrobial properties despite being much shorter in chain-length and lower molecular weight than their natural counterparts. In this work, via computer simulation using coarse-grained model, we have elucidated key mechanisms of membrane-disruption process of a set of isomers of a *β*-peptide, AAK, which is known for their proven antimicrobial activity.

Our ultra-long coarse-grained simulations involving systematically developed coarse-grained model for AAKs, in combination with MARTINI lipid and polarisable water models, were able to capture the spontaneous membrane-disruption processes induced by short-chain *β*-peptide. As a key observation, beyond a threshold peptide/lipid ratio, these coarse-grained simulations revealed formation of spontaneous peptide-induced disordered pore and eventual water-leakage inside the membrane interior. These pores remained stable during the period of 20-40 *µs*-long coarse-grained trajectories. The pore forming peptides are not perfectly vertical with the membrane interface. Rather these peptides showed a non-zero tilt angle with the membrane interface. Interestingly, the pore-forming peptides were found to be considerably extended inside the membrane interior in a ploy for the polar residues to reach out to the headgroups of the distal leaflet. The pore formation propensity increased with increasing P/L ratio, with eventual observation of multiple pores at high P/L ratio. The *β*-peptide-induced pore formation also led to increasing non-uniformity in membrane thickness and disordered lipid tail. Related to this finding, Huang and coworkers(57–59) had majorly used orientational CD spectroscopy to analyze the P/L ratio dependence and pore formation. Their investigation proposed a two-state model of anti-microbial peptides in membrane-parallel and membrane-normal pose. Their analysis based on experimental spectra showed that with increasing P/L ratio, the membrane perpendicular pose is getting prominent. This model is consistent with our current analysis of tilt-angle and water leakage profile as a function of P/L ratio. In particular, the current simulation showed that while at a lower P/L ratio below 1:67, these short-chain beta-peptides remained located at membrane interface at a parallel orientation, with increasing P/L ratio, these beta-peptides reorients themselves into a perpendicular conformation in order to get past the membrane and eventually induce pore formation. These results are in line with Huang group’s proposal. Finally, similar computer simulation using a non globally amphiphilic isomer of the same *β* peptide neither led to any transmembrane pore formation nor induced any water leakage inside the membrane core, which explained its experimentally proven non-potency in antimicrobial activity.

We note that the coarse-grained model used in the current work i.e. MARTINI forcefield together with MARTINI polarisable water is not devoid of criticism. The model has in past been found to require more free energy for pore formation that its all-atom counterpart(60) and has recently been reported(61) for discrepancy in representing the membrane-adsorption process by nanoparticle. However, the employed coarse-grained force field has also been previously successful in prediction of peptide induced pore in the membrane(62, 63). Additionally, the ability of the currently presented coarse-grained model in recovering all-atom simulation results of early stages of peptide-membrane interaction and the observed resilience of the predicted pore in a refined all-atom model (see figure 8) provide confidence on its prediction of long time scale behaviour. The observation of the extended conformation of the *β*-peptides inside the membrane interior in all-atom refined simulation also lends credence to the prediction from the coarse-grained simulation. Nonetheless, setting aside validation of the protein conformational changes by the all-atom simulations, there are also precedent report of larger conformational changes in coarse-grained model (See reference (49) and references of examples there-in). The observation of pore formation in this current coarse-grained simulation is consistent with a previous biochemical investigation of Epand and coworkers(51) in which, a pair of different but closely related 14-helical *β*-peptides were probed for their interaction with model membranes. Using an assay involving large unilamelar vesicles this work had indicated leakage and possibility pore formation and lipid flip-flop by this class of *β*-peptides. The current work provides convincing evidence in support of pore formation by a related but more potent antimicrobial *β*-peptide AAK and brings out the associated molecular mechanism of the full process.

As a major advancement, this work highlights key perspectives of membrane-activity of synthetic biomimetic oligomers. Going beyond precedent computer simulation studies, which have explored mainly membrane-attachment of peptides at a low Peptide/Lipid ratio or which have started with manually preformed specific transmembrane pore(22, 23, 64), our current work takes a step forward and captures the complete process of membrane-disruption via spontaneous formation of barrel-stave pore. The thermodynamic feasibility of membrane disruption via spontaneous pore formation by these rigid 10-residue-long *β*-peptide foldamers were also validated in a wide range of P/L ratios. The superior ability of imparting antimicrobial activity by AAK at a lower MIC than the natural antimicrobial peptides has previously been stressed by Gellman and coworkers(12). In this regard, the observation of transmembrane pore formation via these short-chain peptides and the possibility of chain extension to reach out to the distal leaflet are interesting phenomena. More over the ability to induce trans-membrane pore by AAK at a threshold P/L ratio of 1:67, as found in the current simulation, at considerably lower concentration also speaks of potent membrane-disrupting ability by these *β*-peptides. As we have emphasised earlier in the article, choice of polarisable MARTINI coarse-grained water(28) (in contrast to regular uncharged MARTINI water(27)) was instrumental in capturing the pore formation process by these *β*-peptides, which suggests proper role of water model in capturing the process. The requirement of presence of multiple copies of *β*-peptides for pore formation is a suggestive of cooperative process. In addition, the observation that the reorientation of *β*-peptides from surface-bound state to transmembrane state is facilitated beyond threshold P/L ratios, opens up future direction for systematic quantification via free energy based characterisation of pore formation process(53, 55, 56) using suitable collective variable(65).

The membrane disruption process by antimicrobial agent via pore formation is considered to be a slow process(54, 55, 66), the time scale of which is often prohibitive via all-atom simulation. For example, a 5 *µ*s long all-atom simulation of 10 surface-bound antimicrobial peptides alamethicine did not show any spontaneous membrane insertion.(22) To accelerate insertion of the peptides, the temperature had to be artificially raised to very high temperature and external electric field needed to be employed. In this regard, the current work’s implementation of coarse-grained model in combination with Martini polarisable water is a practical way-forward and more insights can potentially be further harnessed via a resolution-switch to an all-atom model, as have been shown here by the refinement of the coarse-grained model and also in earlier investigation(67). The investigation reported here opens up multitudes of opportunities towards understanding of the mechanistic aspects of the membrane-disruption capability by both natural and biomimetic macromolecules across multiple membrane compositions. Recent works by Gellman and coworkers on exploring random copolymeric biomimetic peptides as cost-effective and efficient antimicrobial agents are being seen as major paradigm shifts(68–70). These works suggest a departure from the preconceived notion of requirement of secondary structure as a key pre-requisite for the manifestation of antimicrobial activity and open up possibilities of venturing into atomistic view of these random copolymeric antimicrobial agents.

## ACKNOWLEDGMENTS

JM thanks Prof. Arun Yethiraj, Prof. Qiang Cui and Prof. Samuel H. Gellman for useful discussions which stimulated the initiation of the project. This work was supported by computing resources obtained from shared facility of TIFR Center for Interdisciplinary Sciences, India. We acknowledge support of the Department of Atomic Energy, Government of India, under Project Identification No. RTI 4007. JM would like to acknowledge research grants received from Ramanujan Fellowship and core research grant provided by the Department of Science and Technology (DST) of India (CRG/2019/001219). PG would like to acknowledge the financial support by INSPIRE Faculty Grant (DST/INSPIRE/04/2015/002495) from DST of India.

## AUTHOR CONTRIBUTION

JM, PG, JK and ND designed the project. JK, DD, ND and JM developed the model. JK, JM and DD performed the coarse-grained simulations. JK and DD performed structural refinement. JK, DD, ND, PG and JM analysed the results. JK and JM wrote the paper.

## SUPPLEMENTAL INFORMATION

Supporting movies of pore formation process at P/L ratio of 1:67, 1:39 and 1:22. All supporting figures referred in this main text (figure S1-S10) are present in the PDF of the supporting information. All gromacs formatted topology and parameter files along with sample configuration for both *β*-peptides investigated in the work can be accessed from the following Github link: https://github.com/JMLab-tifrh/coarse-grained-models-of-beta-peptides

