## Supplemental figure and tables for "Spontaneous Transmembrane Pore Formation by Short-chain Synthetic Peptide"

### Description of Movies

- Movie S1: The trajectory capturing the trans-membrane pore formation process by GA isomer at a P/L ratio of 1:67 ( 5 copies of peptides)
- Movie S2: The trajectory capturing the trans-membrane pore formation process by GA isomer at a P/L ratio of 1:38 ( 10 copies of peptides)
- Movie S3: The trajectory capturing the trans-membrane pore formation process by GA isomer at a P/L ratio of 1:22 ( 15 copies of peptides)

  Color code used in all movies: Color code: tan : Phosphate groups of lipids, orange : Plolarized water, blue :: Tyrosin, green : Lysin, red : ACHC residues.

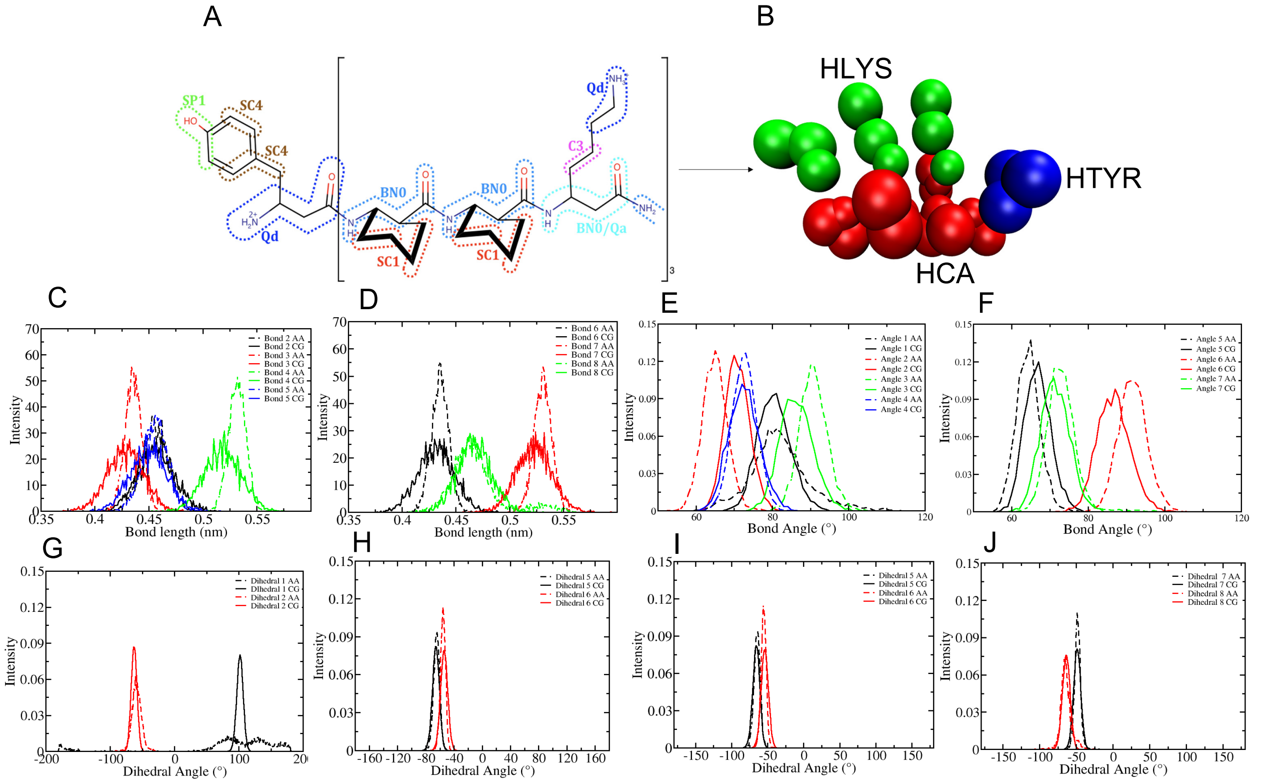


Figure S1: Comparison of distributions of bonded terms for coarse-grained model of GA isomer with that of all-atom model.

Table S1: Details of the systems for simulation using non-GA isomer of AAK. All $\beta$peptide copies were initially placed in aqueous phase close to one side of the membrane. Each system was charge-neutralised by adding necessary counter ions. As a common point, the water layer was thicker in system involving non GA isomers, to avoid them artificially reaching out to other leaflet due to periodic boundary condition.

| System | Number of water | P/L ratio | Simulation Time($\mu$s) $\times$ number of replica |
|---|---|---|---|
| $\beta$peptide-less bilayer | 6538 | 0 | 5 $\mu$s $\times$ 2 |
| 1 non-GA in 388 lipid | 7381 | 1:388 | 20 $\mu$s $\times$ 2 |
| 5 non-GA in 336 lipid | 8916 | 1:89 | 40 $\mu$s $\times$ 4 |
| 10 non-GA in 388 lipid | 11854 | 1:39 | 20 $\mu$s $\times$ 4 |
| 15 non-GA in 336 lipid | 10236 | 1:22 | 20 $\mu$s $\times$ 4 |

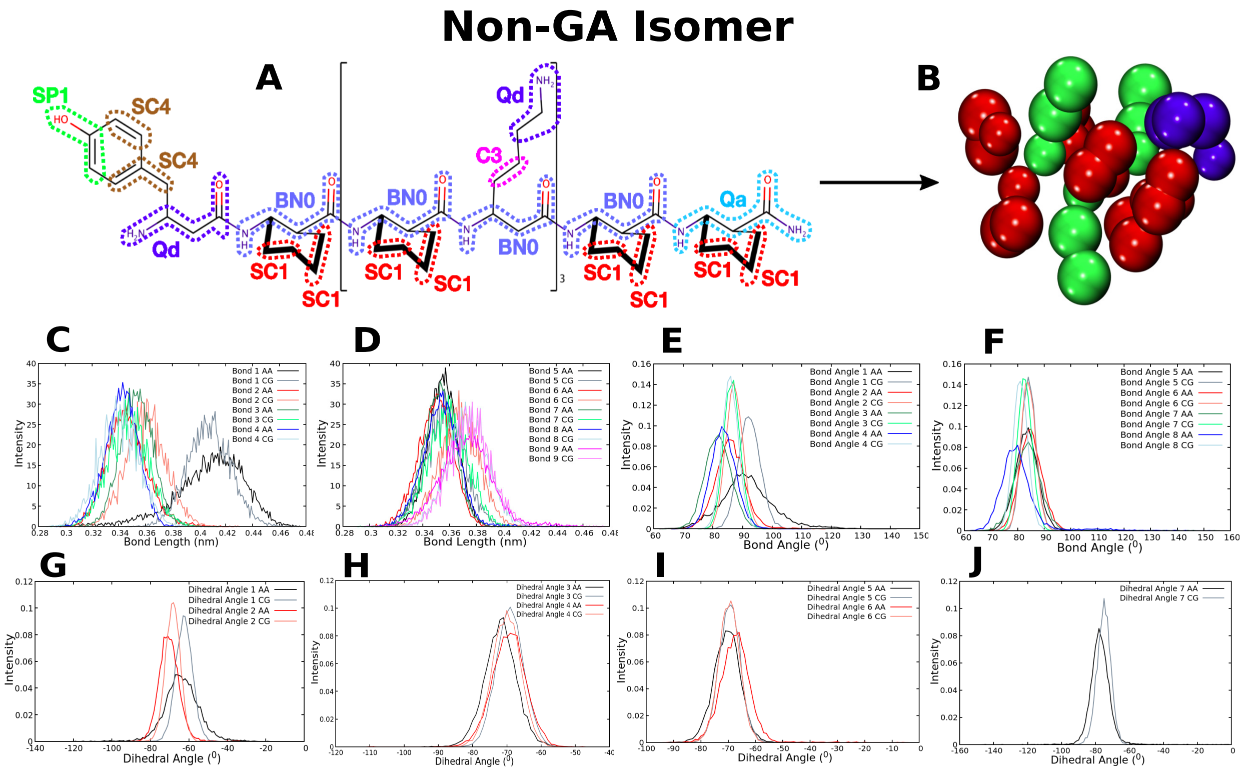


Figure S2: Mapping scheme and comparison of distributions of bonded terms for coarse-grained model of non-GA isomer with that of all-atom model.

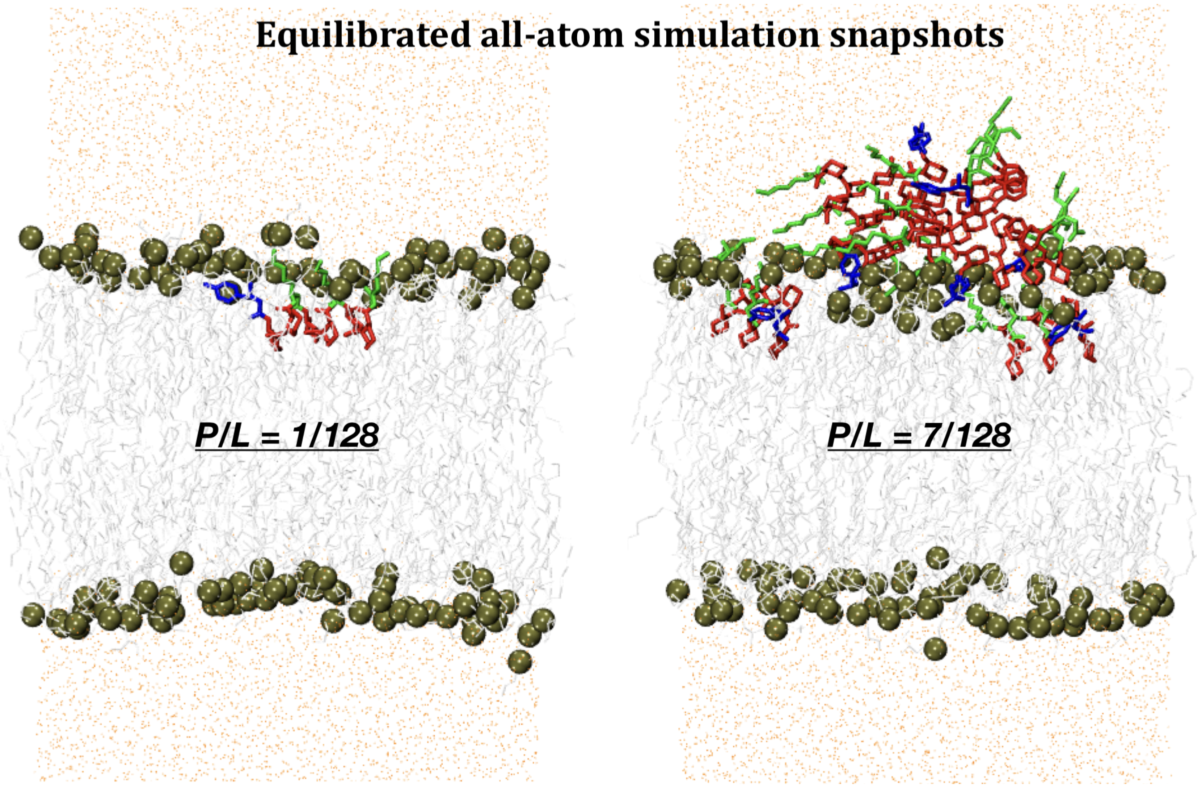


Figure S3: All-atom results with single and multiple $\beta$-peptides

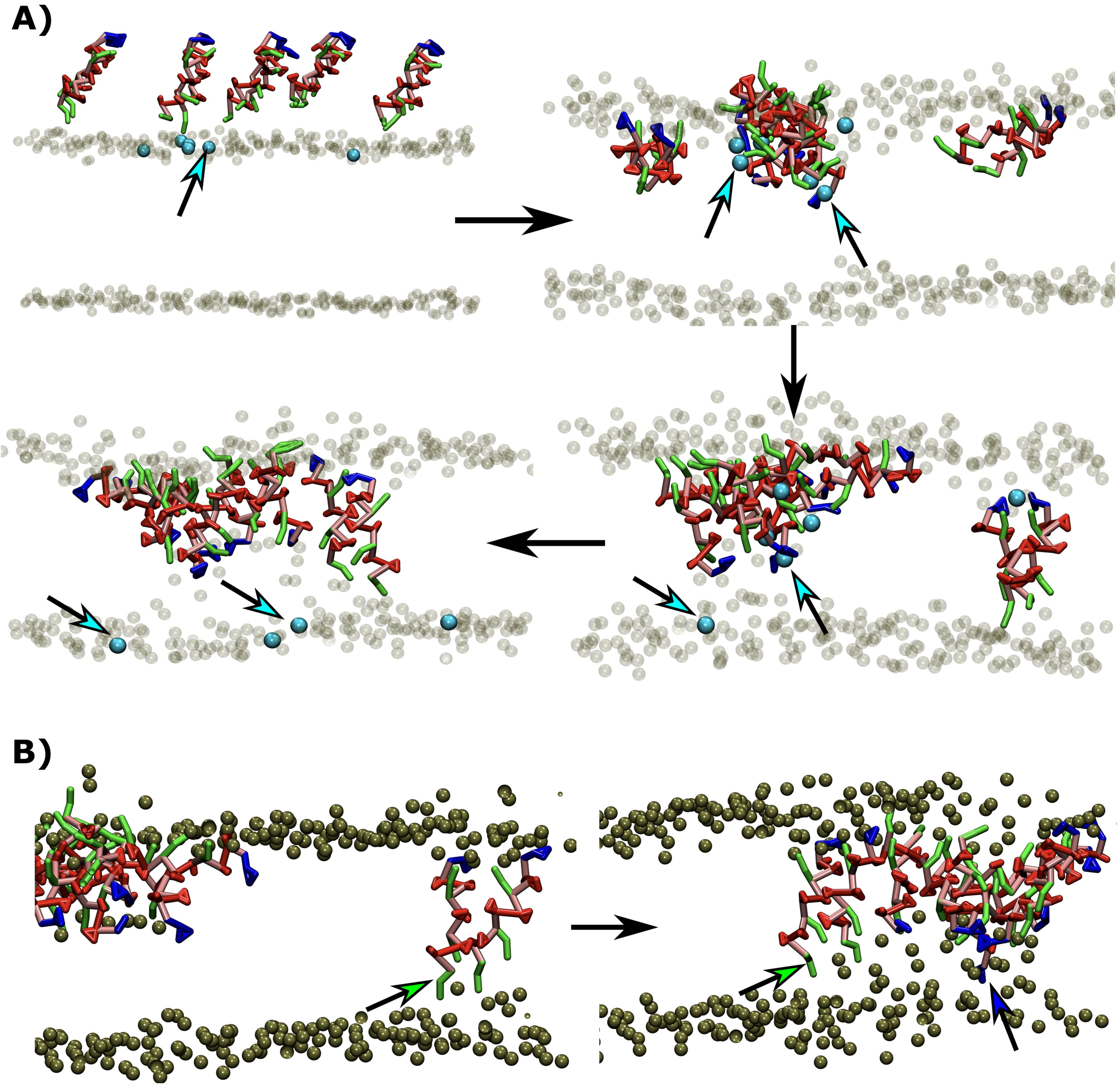


Figure S4: A. Zoomed-in view of early steps of peptide-induced pore formation, specifically highlighting the displacement of lipid head groups (cyan colors) by peptide-aggregate. B. Snapshot indicating that the polar and charged residues extend themselves to the distal head groups of bilayer

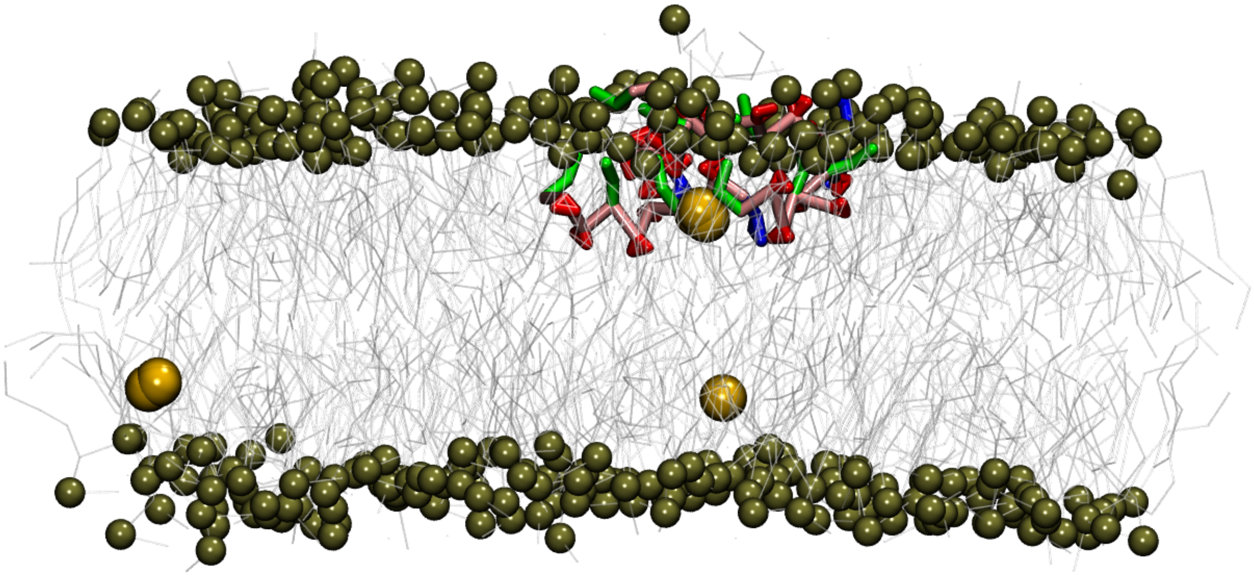

Figure S5: Representative snapshot of final configuration at P/L ratio less than 1:67 (involving 4 peptides in 336 lipids) in coarse-grained simulation.

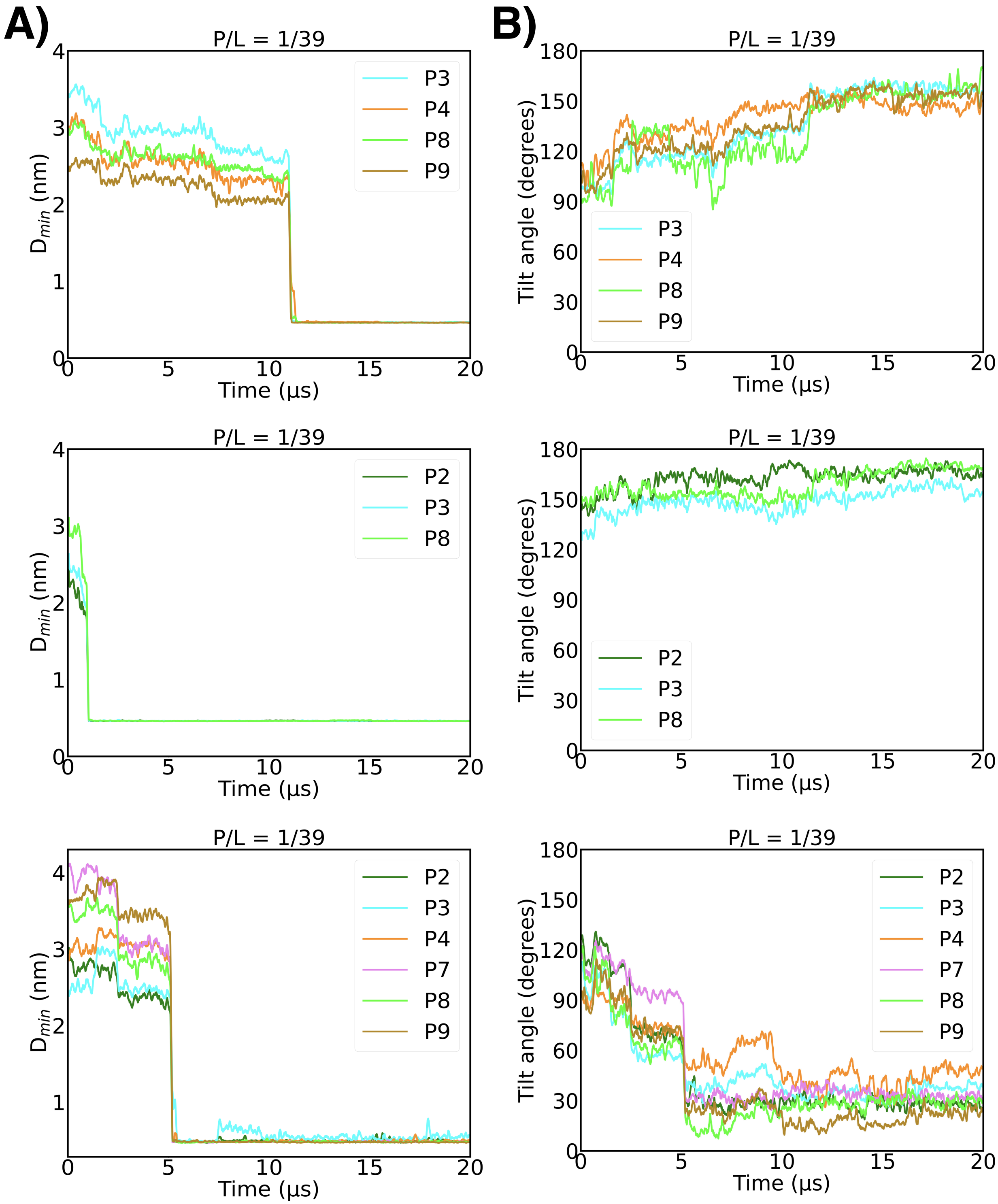


Figure S6: A. Time profile of extent of peptide insertions in multiple independently performed trajectories for P/L of 1:38. Here, minimum distance between the phosphate group of distal leaflet and the peptide ($d_{min}$) has been considered as the metric. The sudden drop in the value of $d_{min}$ signifies the onset of peptide internalisation and pore formation. Only the time profile of pore-forming peptide has been shown. B. Time profile of tilt angle of the peptides with the membrane interface at each of the respective trajectories.

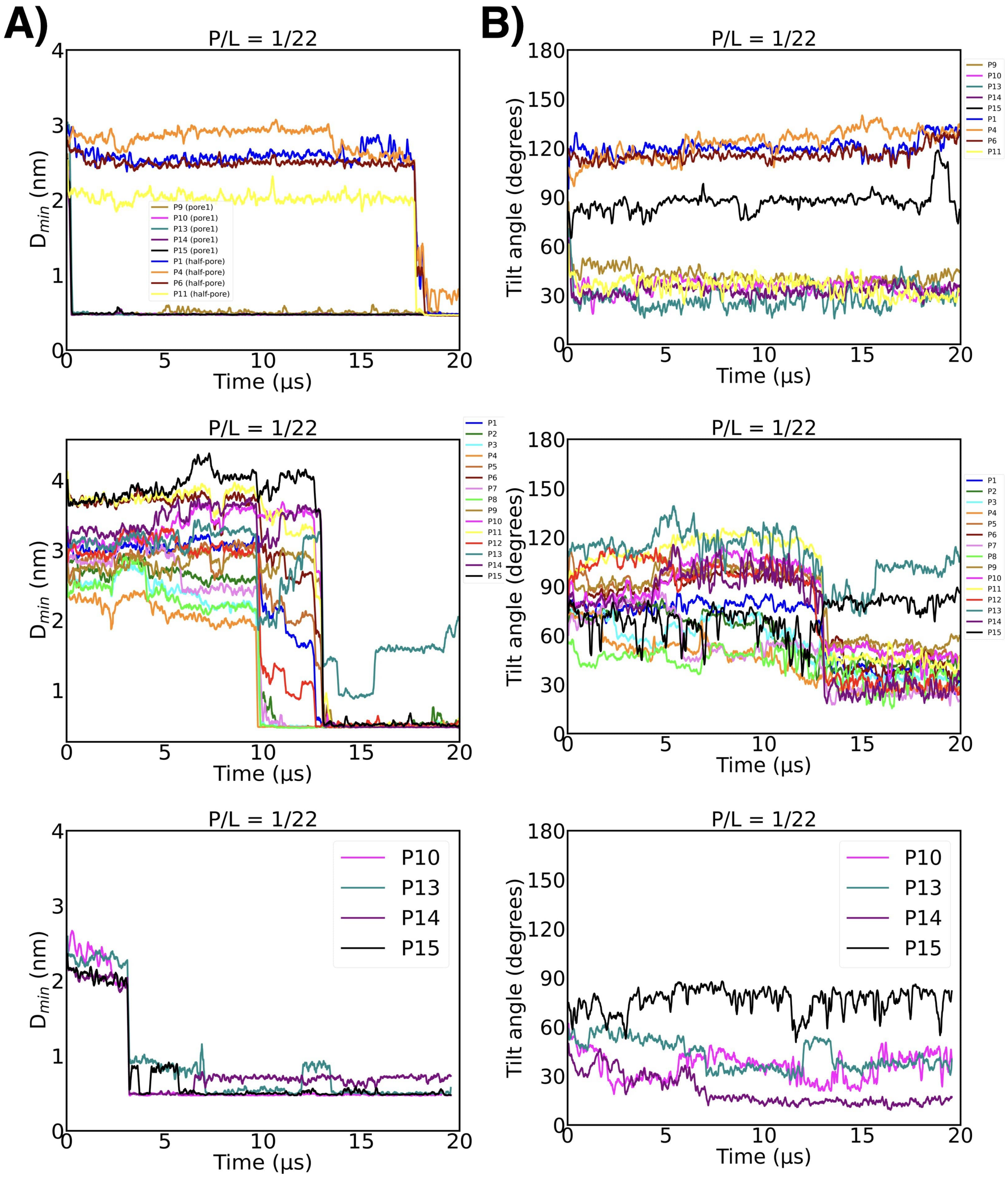


Figure S7: A. Time profile of extent of peptide insertions in multiple independently performed trajectories for P/L of 1:22. Here, minimum distance between the phosphate group of distal leaflet and the peptide ($d_{min}$) has been considered as the metric. The sudden drop in the value of $d_{min}$ signifies the onset of peptide internalisation and pore formation. Only the time profile of pore-forming peptides (annotated by 'P1, P2...') has been shown. B. Time profile of tilt angle of the peptides with the membrane interface at each of the respective trajectories.

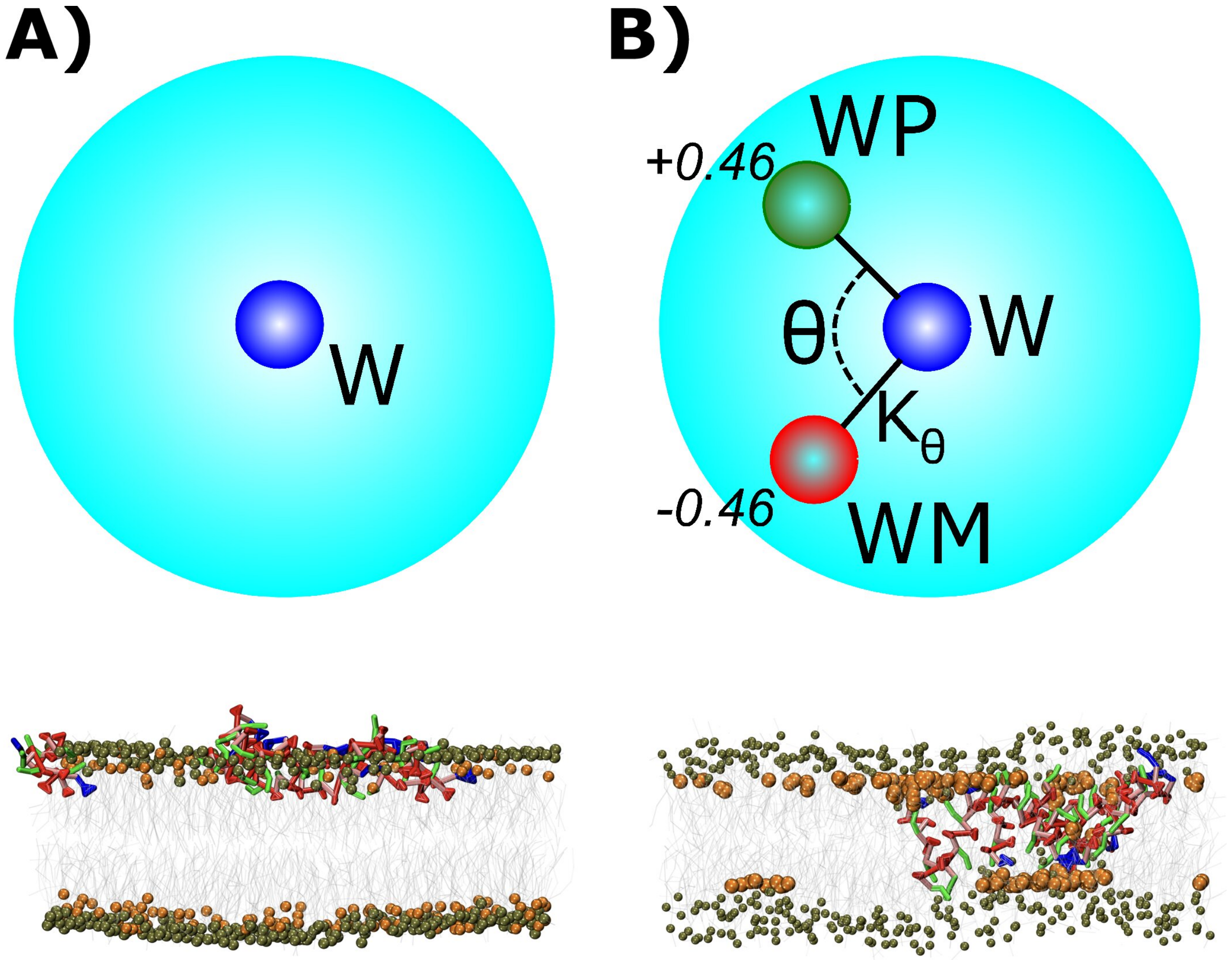


Figure S8: Comparison of Representative snapshot of final configuration at P/L ratio 1:39 in which A) regular LJ coarse-grained MARTINI water has been used.B) polarisable MARTINI water has been used.

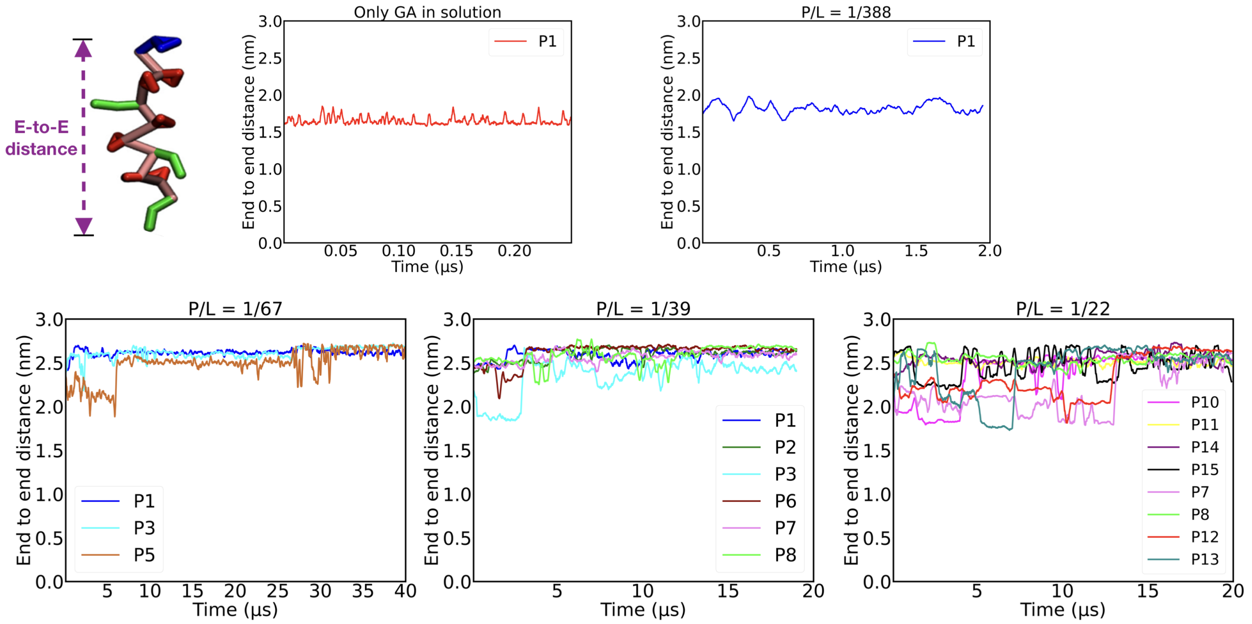


Figure S9: The time profile of end-to-end distance of individual $\beta$-peptides (GA isomers) which are constituent of pore. Shown for all P/L ratios studied here. The increase in the value of end-to-end distance at higher P/L ratios is evident.

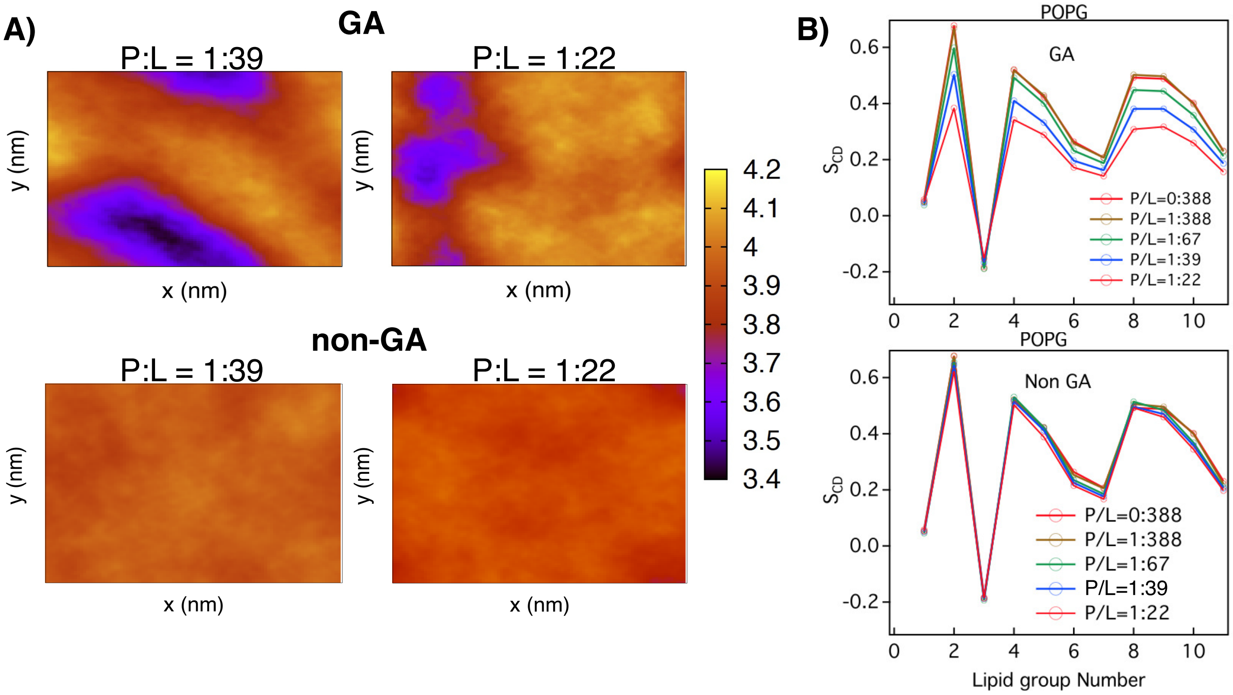


Figure S10: A. Comparison of bilayer thickness profile between GA and non GA isomers of AAK at two P/L ratios. B. Comparison of POPG lipid order parameter between GA and non GA isomers .
